# SYNAPTIC PLASTICITY IN THE INJURED BRAIN DEPENDS ON THE TEMPORAL PATTERN OF STIMULATION

**DOI:** 10.1101/2023.12.13.571587

**Authors:** Quentin S. Fischer, Djanenkhodja Kalikulov, Gonzalo Viana Di Prisco, Carrie A. Williams, Philip R. Baldwin, Michael J. Friedlander

**Author notes:** Address correspondence to: Dr. Michael J. Friedlander, Fralin Biomedical Research Institute at VTC, 2 Riverside Circle, Roanoke VA 24016 USA. Department of Pharmacology and Toxicology, Indiana University School of Medicine, 320 W. 15th St., NB-400, Indianapolis, IN 46202 USA. Department of Biochemistry and Molecular Pharmacology, Baylor College of Medicine, 1 Baylor Plaza, Houston TX 77030 USA.

## Abstract

Neurostimulation protocols are increasingly used as therapeutic interventions, including for brain injury. In addition to the direct activation of neurons, these stimulation protocols are also likely to have downstream effects on those neurons’ synaptic outputs. It is well known that alterations in the strength of synaptic connections (long-term potentiation, LTP; long-term depression, LTD) are sensitive to the frequency of stimulation used for induction, however little is known about the contribution of the temporal pattern of stimulation to the downstream synaptic plasticity that may be induced by neurostimulation in the injured brain. We explored interactions of the temporal pattern and frequency of neurostimulation in the normal cerebral cortex and after mild traumatic brain injury (mTBI), to inform therapies to strengthen or weaken neural circuits in injured brains, as well as to better understand the role of these factors in normal brain plasticity. Whole-cell (WC) patch-clamp recordings of evoked postsynaptic potentials (PSPs) in individual neurons, as well as field potential (FP) recordings, were made from layer 2/3 of visual cortex in response to stimulation of layer 4, in acute slices from control (naïve), sham operated, and mTBI rats. We compared synaptic plasticity induced by different stimulation protocols, each consisting of a specific frequency (1 Hz, 10 Hz, or 100 Hz), continuity (continuous or discontinuous), and temporal pattern (perfectly regular, slightly irregular, or highly irregular). At the individual neuron level, dramatic differences in plasticity outcome occurred when the highly irregular stimulation protocol was used at 1 Hz or 10 Hz, producing an overall LTD in controls and shams, but a robust overall LTP after mTBI. Consistent with the individual neuron results, the plasticity outcomes for simultaneous FP recordings were similar, indicative of our results generalizing to a larger scale synaptic network than can be sampled by individual WC recordings alone. In addition to the differences in plasticity outcome between control (naïve or sham) and injured brains, the dynamics of the changes in synaptic responses that developed during stimulation were predictive of the final plasticity outcome. Our results demonstrate that the temporal pattern of stimulation plays a role in the polarity and magnitude of synaptic plasticity induced in the cerebral cortex while highlighting differences between normal and injured brain responses. Moreover, these results may be useful for optimization of neurostimulation therapies to treat mTBI and other brain disorders, in addition to providing new insights into downstream plasticity signaling mechanisms in the normal brain.

## INTRODUCTION

A wide variety of neurological and psychiatric effects result from brain injuries like mTBI.^1–6^ The capacity of the brain to adapt to injury and undergo restorative plasticity throughout the lifespan is well-documented.^7–14^ Up- and down-regulation of the strength of synaptic transmission such as LTP and LTD represent processes implicated as biological substrates of learning and memory.^15–27^ Such changes have been observed throughout the brain including the: hippocampus,^28–31^ visual cortex,^32–34^ olfactory cortex,^35,36^ somatosensory and motor cortices,^37–39^ striatum,^40,41^ and cerebellum.^42,43^ LTP and LTD are also implicated in functional reorganization after brain injury,^44–49^ and during neurorehabilitation for brain injury or disease.^50–56^ It is of interest to learn whether these processes may be selectively evoked to facilitate a compensatory rebalancing of the relative strengths of particular synaptic pathways after mTBI and thus become a therapeutic target for restoring functionality.^57–59^ Moreover, with the availability of various neurostimulation technologies such as: deep brain stimulation,^60,61^ transcranial focused ultrasound,^62,63^ transcranial magnetic stimulation,^64–66^ transcranial direct current stimulation,^67,68^ and more,^69^ to treat conditions such as Parkinson’s disease,^70–72^ Alzheimer’s disease,^73–75^ seizure disorders,^76–78^ and major depression,^79–82^ there is value in better understanding the effects of these interventions on synaptic plasticity in a well-controlled experimental environment.

Although, some clinical uses of deep brain stimulation have begun to take into account temporal pattern,^83–85^ most experimental approaches to studying the induction mechanisms for LTP and LTD hold temporal patterns regular and constant, with a brief period of high frequency afferent stimulation,^86,87^ a protracted period of low frequency presynaptic stimulation,^31,88^ or a series of conjunctions of temporally coincident pre- and post-synaptic activation.^89–91^ While varying stimulation frequency, these protocols maintain a regular interval between each stimulus (i.e. all interstimulus intervals, ISIs, are equal). These temporally regular epochs of synaptic stimulation have been widely applied in isolated *in vitro* experimental preparations where the neurons are usually quiescent until stimulated. However, *in vivo* many neurons exhibit ongoing activity, continuously receiving synaptic inputs, many of which are evoked by presynaptic action potentials.^92–96^ Moreover, the temporal pattern of action potentials and subsequent evoked synaptic activity is often highly irregular with varying distributions of ISIs.^97–102^ Thus, it is important to understand the contribution of temporal pattern (vs. frequency alone) of evoked synaptic activity *per se* to the induction of synaptic plasticity in the normal brain and whether there are different temporal pattern requirements for accessing downstream synaptic plasticity signaling cascades after mTBI.

Since the neocortex is often subjected to mTBI and the synaptic circuitry of the visual cortex is well-characterized, we chose this area to evaluate the role of the temporal pattern of evoked synaptic activity. In particular, we were interested to learn whether the major synaptic drive from the neurons of the primary thalamic input zone (layer 4, L4), onto the layer 2/3 (L 2/3) pyramidal neurons that transmit information over long distances between cortical areas, react differently to variations in the temporal pattern of conditioning inputs in the normal brain and after mTBI. We focused on three stimulation frequencies known to play a role in plasticity in neocortex (1, 10, 100 Hz), three temporal patterns (perfectly regular: all ISIs are identical with a coefficient of variation or CV = 0.0, slightly irregular: Gaussian distribution of ISIs with a CV = 0.2, and highly irregular: Poisson distribution of ISIs with a CV = 1.0), and two types of conditioning epochs (continuous or discontinuous).

## METHODS (Details in Supplementary Materials)

### Animal use and experimental design

All procedures were approved by the Institutional Animal Care and Use Committees of Virginia Tech or Baylor College of Medicine and followed the National Research Council’s *Guide for the Care and Use of Laboratory Animals*. At 8-9 weeks of age, male Long-Evans rats were single-housed and assigned to one of three treatment groups: control (naïve), sham (sham operated), or mTBI (controlled cortical impact, CCI). Two to three weeks later these animals were used in: righting reflex, electrophysiology, and/or calcium imaging experiments.

### Mild traumatic brain injury model

At 8-9 weeks of age the mTBI group received a CCI using a well-established model.^103–105^ Surgery was performed under isoflurane anesthesia (1–4%). Animals were placed on a homeostatically controlled heating plate and mounted in a stereotaxic frame. The scalp was shaved, cleaned with alcohol and betadine, and incised to expose the skull. A 7 mm diameter craniotomy was made over the right parietal cortex immediately posterior to bregma and lateral to midline. A direct lateral CCI (2.5 mm depth, 3.0 m/s velocity, 100 ms duration, angled at 45° from vertical) was delivered through intact meninges using a Benchmark stereotaxic impactor (Leica) with a 6 mm diameter spherical head. After the CCI, we administered a long-lasting analgesic (buprenorphine SR, 1.2 mg/kg SC; ZooPharm), the craniotomy and incision were repaired, anesthesia was discontinued, and the rat monitored until fully recovered. Shams received identical treatment except for the CCI.

### Righting reflex

At 10-12 weeks of age (2-3 weeks after treatment: control, sham, or mTBI) the first cohort of 45 rats was examined to assess righting latency. Individuals were placed in an induction chamber and anesthetized with isoflurane (4% for 120 s). The rat was removed, placed in a supine position and righting latency (the time required for the rat to right itself to an upright posture standing on all four paws) was measured.

### Brain slice preparation

At 10-12 weeks of age (2-3 weeks after treatment: control, sham, or mTBI) a second cohort of 389 rats was anesthetized with ketamine/xylazine (75 and 10 mg/kg, IM) then decapitated. The brain was removed and placed in ice-cold cutting solution containing (in mM): 124 NaCl, 2 KCl, 3.5 MgCl_2_, 0.2 CaCl_2_, 1.25 KH_2_PO_4_, 26 NaHCO_3_, 11 dextrose, with pH 7.4 and saturated with carbogen (95% O_2_ + 5% CO_2_). Coronal slices (300 µm) were prepared from the right visual cortex (ipsilateral to the mTBI) and placed in artificial cerebral spinal fluid (ACSF—containing in mM: 124 NaCl, 2 KCl, 2 MgCl_2_, 2 CaCl_2_, 1.25 KH2PO_4_, 26 NaHCO_3_, 11 dextrose, with pH 7.4 and saturated with carbogen). Slices were incubated in ACSF at 33 ± 1°C for 20 min and then at room temperature for > 40 min prior to recording.

### *In vitro* electrophysiology

For recording, slices were transferred to a submersion chamber perfused with 3 ml/min of ACSF at 33±1°C. A bipolar stimulating electrode (FHC #30213) was placed in L4. Glass micropipettes (OD 1.5 mm, ID 1.12 mm; A-M Systems) were pulled on a vertical puller (PC-10; Narishige) to make FP pipettes (1 ± 0.5 MΩ), or WC patch pipettes (4 ± 1 MΩ). FP pipettes were filled with ACSF and placed in L2/3. WC patch pipettes were filled with pipette solution (containing in mM: 115 K-gluconate, 20 KCl, 10 Hepes, 4 NaCl, 4 Mg-ATP, 0.3 Na-GTP, and 4 phosphocreatine-Na; pH 7.4; 280-290 mOsm) and placed in L2/3 (∼100 µm deep to the FP pipette). Biocytin (0.5%; Sigma-Aldrich) added to the pipette solution allowed post-hoc identification of patched cells (only pyramidal cells located in L2/3 were analyzed).

Recordings were made using a MultiClamp 700B amplifier, digitized at 20 kHz with a Digidata 1440A, and collected and analyzed with pCLAMP 10 software (Axon Instruments, Molecular Devices). Recordings of evoked PSP and FP amplitudes were measured as the difference between baseline and peak. Recordings with unstable pre-conditioning response amplitudes, membrane potentials more positive than -60 mV, high access resistance (> 40 MΩ, or > 20% of the membrane resistance for that cell), or a > 20% decrease in input resistance were discarded. Intrinsic firing properties of patched cells were tested by injecting a 100 ms depolarizing current pulse and cells that did not exhibit regular spiking typical of excitatory cells^106,107^ were discarded. Only one recording was made from each slice, so that a single stimulation protocol was applied in each case.

### Stimulation protocols

Afferent stimulation consisted of constant current square waves (100 µs, 30-500 µA; set to evoke ∼40% of the maximal PSP response) output by a stimulus isolation unit (SIU; A365, World Precision Instruments). Stimulation was optimized to produce PSPs, but in some cases (∼30%) also evoked FPs which were recorded simultaneously. Pre-conditioning baseline recordings of evoked PSP (and FP) peak amplitudes were made at 0.1 Hz for 10 min. This was followed by a 90 or 900 s period of synaptic conditioning, consisting of 900 pulses at 1, 10, or 100 Hz. Conditioning was either continuous (1 Hz for 900 s, or 10 Hz for 90 s; Fig. 1A-B) or discontinuous (9 epochs of 100 pulses each at 10 or 100 Hz, separated by 8 equal rest intervals consisting of probe trials at 0.1 Hz, with a total duration of 900 s; Fig. 1C). We included the discontinuous protocols since continuous stimulation at high frequency can induce presynaptic fatigue that might interact with long-term synaptic plasticity.^108,109^ Post-conditioning evoked PSP (and FP) peak amplitudes were evaluated at 0.1 Hz for 20 min. Conditioning stimulation was delivered with one of three temporal patterns: perfectly regular (CV 0.0; Fig. 1D-E), slightly irregular (CV 0.2; Fig. 1F-G), or highly irregular (CV 1.0; Fig. 1H-I) each with the same mean frequency. Conditioning stimulation was generated using MatLab (MathWorks), transformed with a digital analog converter (BNC 2090A, National Instruments), triggered with a pulse stimulator (Master-9, A.M.P.I.), and output via the SIU, giving us precise control of frequency, continuity and temporal pattern.

**Fig. 1.**
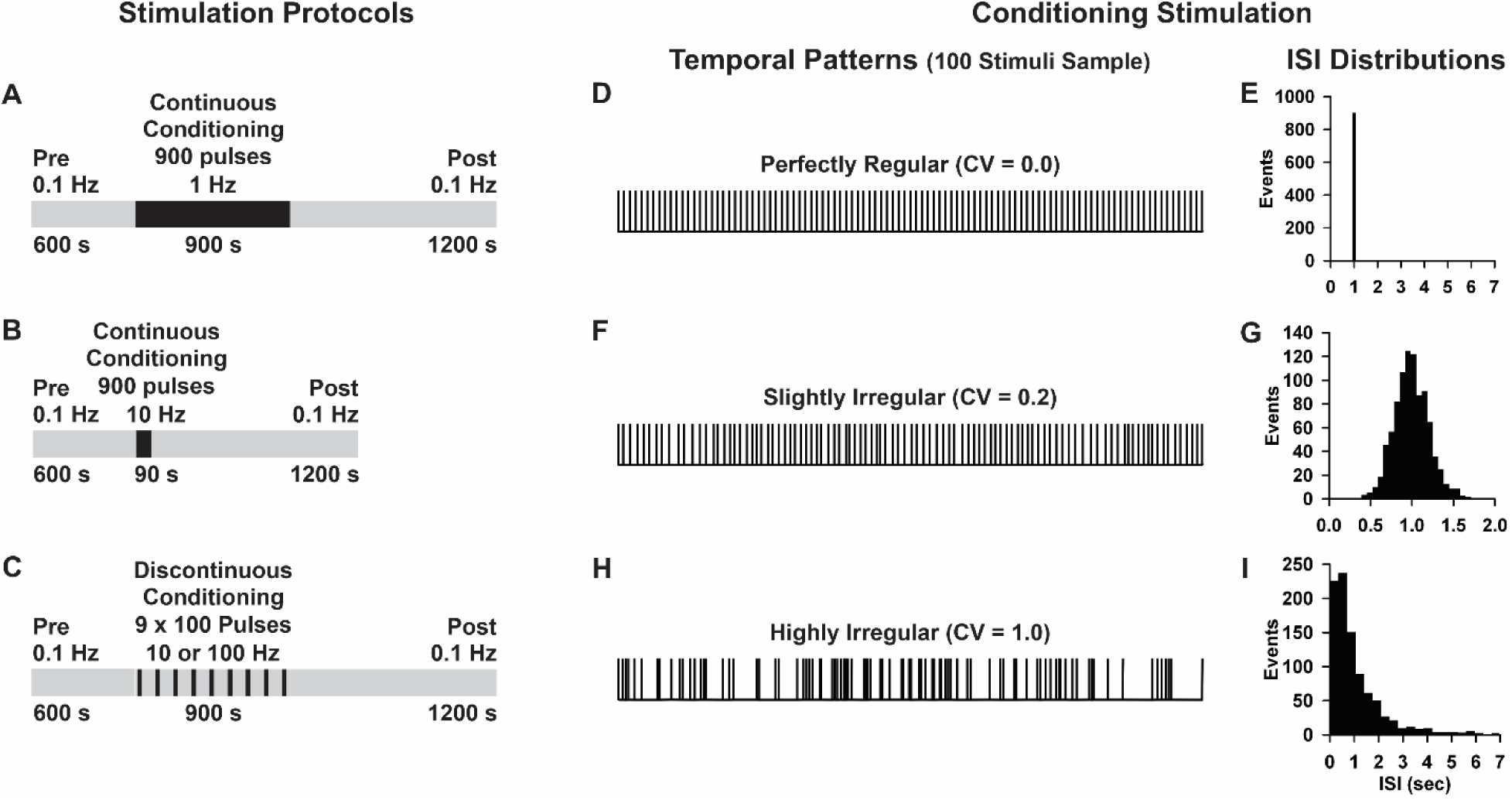
Stimulation protocols, temporal patterns, and interstimulus interval distributions. (**A-C**) Stimulation protocols for: 1 Hz continuous, 10 Hz continuous, and 10 Hz (or 100 Hz) discontinuous conditioning, respectively. Grey bands represent pre- and post-conditioning periods, while black bands represent intervals of conditioning stimulation. Note that for the discontinuous stimulation protocols shown in **C** the conditioning period consists of alternating stimulation epochs (black bands) and rest intervals (grey bands). Rest intervals (10 Hz = 101.25 sec each, 100 Hz = 111.372 sec each) have probe trials at 0.1 Hz. (**D-E**) Example of perfectly regular conditioning **D**, with all ISIs equal **E**. (**F-G**) Example of slightly irregular conditioning **F**, where ISIs have a Gaussian distribution **G**. (**H-I**) Example of highly irregular conditioning **H**, where ISIs have a Poisson distribution **I**. Note that the times shown in panels **E**, **G**, and **I** are based on 1 Hz stimulation (10 Hz or 100 Hz stimulation are identical in pattern, but 10 or 100 times shorter in duration, respectively).

### Temporal patterns of conditioning stimulation

Conditioning stimulation temporal patterns were generated by creating ISIs according to the family of distributions given by:

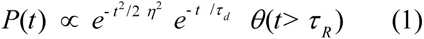

where τ_R_ is the refractory period, and η (>0) and τ_d_ are temporal parameters that govern the single peak of the distribution. For a perfectly regular distribution, we had -τ_d_/η >> 1, and for a highly irregular distribution η/τ_d_ >> 1. We solved for the parameters η and τ_d_, to yield a desired mean, m, and standard deviation (std) or coefficient of variation (CV):

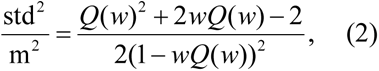

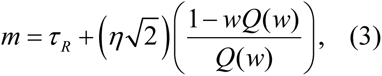

where 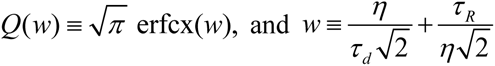. Since equations (2) and (3) are monotonic, one can invert the equations to find η and τ_d_ in terms of m and std. Thus, it was possible to change a set of ISIs at one CV to a related set of ISIs at another CV. The cumulative distribution function corresponding to the event probability distribution function given by equation (1) above is:

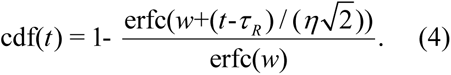

We randomly assigned numbers drawn uniformly on the unit interval to ISIs in the following way. For each random number, r, we inverted equation (4) to find a *t* that represented an ISI affiliated with a given mean and CV. By specifying a different mean and CV (and therefore different η and τ_d_, or equivalently, different η and *w*) in equation (4) we derived related sets of ISIs, using a single set of random numbers.

### Analyzing electrophysiology data

We measured evoked PSP (or FP) peak amplitudes during the last 5 min of pre- and post-conditioning (using pCLAMP) and calculated a post/pre ratio. We evaluated the effects of mTBI, as well as stimulus frequency, temporal pattern, and continuity on the induction of synaptic plasticity [defined as: LTD, a significant (p < 0.05, *t-test*) decrease (≥ 10%) in post/pre ratios; LTP, a significant increase (≥ 10%) in post/pre ratios; or no change (NC), post/pre ratios that changed < 10% and/or were not significantly different]. For these same time periods (the last 5 min of pre- and post-conditioning), we used pCLAMP to calculate the average PSP: half width (at half height), rise time (10-90%) and decay time (90-10%); as well as the membrane potential and input resistance for each cell.

We also analyzed the kinetic profiles of conditioning responses for individual recordings in the groups that showed a significant difference between control and mTBI. For continuous stimulation protocols, we took all data points generated during the conditioning period and calculated exponential and linear best fits, from which we determined the slope. For discontinuous stimulation protocols, best fits were calculated for each of the nine conditioning sub-epochs and the slopes averaged. Further, we calculated the mean of all conditioning response amplitudes (average) as well as the average of the initial 10% (initial) and final 10% (final). For all recordings within a given stimulation protocol, a linear regression analysis was performed to determine the degree to which each factor (exponential slope, linear slope, average amplitude, initial amplitude and final amplitude) was correlated with plasticity outcome (post/pre ratio).

### Calcium imaging

In a subset of the WC patch experiments, the ratiometric calcium sensitive dye Fura-4F (100 µM, Thermo Fisher Scientific) was added to the pipette solution. Baseline somatic calcium concentrations were calculated as the ratio of bound to unbound calcium minus background (AxioVision; Zeiss) during the last five minutes of pre-conditioning.

### Statistics

We used SigmaPlot (Systat Software) to determine statistical significance except where noted. Normality was assessed with a Shapiro-Wilk test. For normally distributed data, we used a *t-*test, or an ANOVA followed by Fisher’s least significant difference method for multiple comparisons (Fischer’s LSD). For datasets with non-normal distributions, we used a Mann-Whitney U test (MWU), or a Kruskal-Wallis one-way ANOVA on ranks (Kruskal-Wallis) followed by Dunn’s method for multiple comparisons with Sidak correction (Dunn’s corrected; statskingdom.com/kruskal-wallis-calculator.html). Group data are presented as the mean ± SEM. To assess whether specific characteristics of conditioning responses were predictive of plasticity outcome, for each factor, we calculated a correlation coefficient and determined significance using linear regression. We considered differences statistically significant at *p* < 0.05 (except for Dunn’s corrected which was considered significant at *p* < 0.017).

## RESULTS

### Efficacy of our mTBI model

To establish the efficacy of our CCI model for mTBI we used several independent methods. First, we used a behavioral measure, recovery of righting reflex, to assess motor and cognitive function in control, sham and mTBI rats. Righting latency was significantly impacted by treatment group (*p* = 0.04), and was longer in mTBI than in control (*p* = 0.04) or sham (*p* = 0.02; Fig. 2A) groups. Note that for this and all subsequent figures we show control data in black, sham data in blue, and mTBI data in red. Second, we measured intracellular calcium concentration in a subset of the L2/3 pyramidal neurons used for our WC patch-clamp recordings. Intracellular calcium concentrations were significantly different between treatment groups (*p* < 0.001), and were higher in the mTBI than in the control (p < 0.001) or sham group (p < 0.001; Fig. 2B). Third, we measured the intrinsic properties of this same subset of cells, and found that input resistance was significantly affected by treatment group (*p* = 0.005), being higher in mTBI than in control (*p* = 0.001) or sham (*p* = 0.008; Fig. 2C) groups; while membrane potential did not differ between control, sham, or mTBI groups (*p* = 0.38; Fig. 2D).

**Fig. 2.**
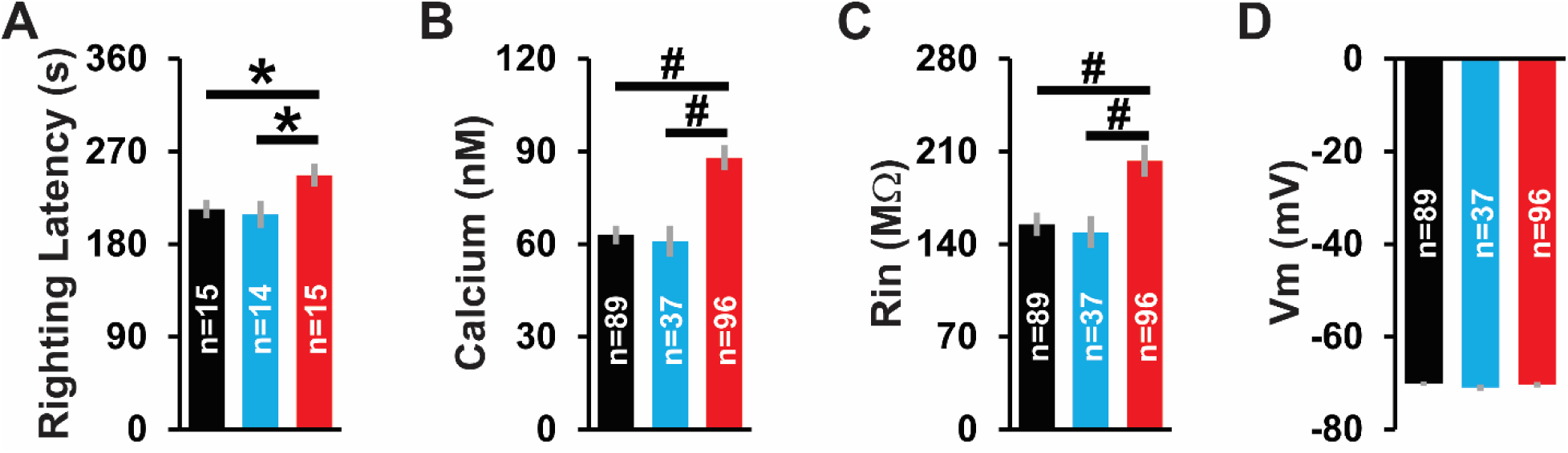
Efficacy of our CCI model for mTBI. (**A**) Righting latency, (**B**) intracellular calcium, and (**C**) input resistance are significantly increased in mTBI verses control or sham groups. The *p* values and statistical tests are: *p* = 0.04 ANOVA, *p* < 0.001 Kruskal-Wallis, *p* = 0.005 Kruskal-Wallis, respectively. Subsequent pairwise comparisons show significant differences: \**p* < 0.05 Fisher’s LSD, #*p* < 0.017 Dunn’s corrected. For this and all subsequent figures we show control data in black, sham data in blue and mTBI data in red. Values are mean ± SEM, n = number of observations/group. (**D**) Membrane potential did not differ between treatment groups (control, sham, mTBI; *p* = 0.38 Kruskal-Wallis).

### Effect of temporal pattern and treatment condition on plasticity outcome

WC patch-clamp experiments were conducted to investigate the role of temporal pattern on synaptic plasticity in control, sham and mTBI groups. Figure 3 shows example recordings of evoked PSPs and time plots of individual recordings in response to 1 Hz continuous highly irregular conditioning (*cf.* Fig. 1A, 1H-I). Note the substantial decrease in post-conditioning PSP amplitude, indicative of LTD, in control and sham examples (Fig. 3A-B). In contrast, there was a substantial increase in post-conditioning PSP amplitude, indicative of LTP, in the mTBI example (Fig. 3C).

**Fig. 3.**
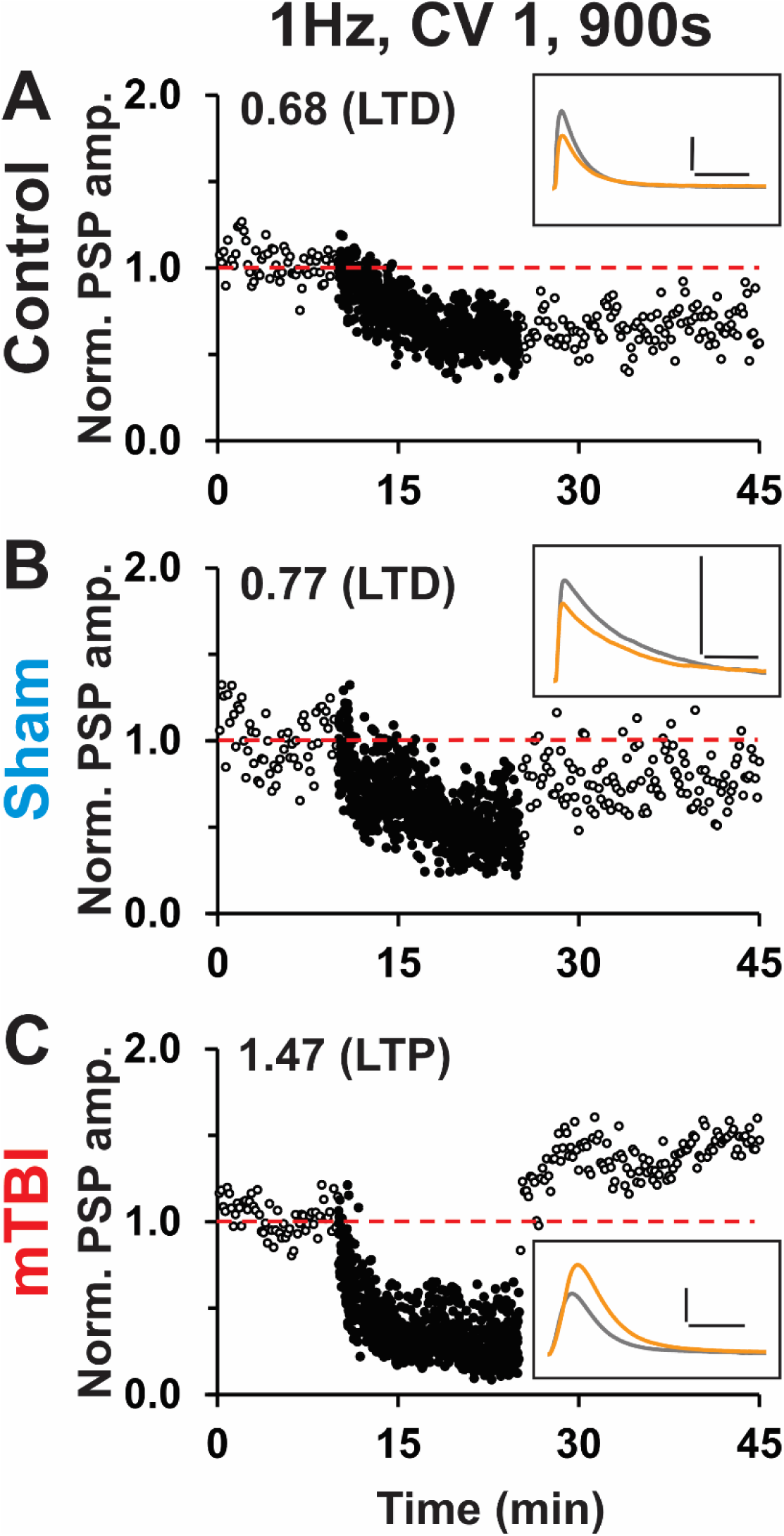
Individual time plots of WC responses to highly irregular 1 Hz continuous conditioning. (**A-C**) Representative examples from: control, sham, and mTBI groups. Black open circles = pre-/post-conditioning responses, black filled circles = conditioning responses, red dotted line = baseline. Numerical values = post/pre ratios and corresponding plasticity outcome (LTD, NC or LTP). Insets: PSP waveforms averaged over the last 5 min of pre-(grey) or post- (orange) conditioning (scale bars: 5 mV, 50 ms).

Grouped results for our WC recording experiments for the various frequencies, temporal patterns, and treatment groups are shown in subsequent figures. Continuous conditioning at 1 Hz induced LTD in both control groups (Fig. 4A-C) and mTBI groups (Fig. 4D-E) in all cases except one—highly irregular conditioning (CV 1.0) in mTBI rats resulted in LTP (Fig. 4F). Temporal pattern (CV 0.0, CV 0.2, CV 1.0) had no effect on plasticity outcome in control groups (*p* = 0.50; Fig. 4A-C, 4G). However, for mTBI groups, there was a significant difference in plasticity outcome between the three temporal patterns (*p* = 0.002; Fig. 4D-F, 4H) with highly irregular conditioning (CV 1.0) showing significant potentiation (verses: CV 0.0 *p* = 0.007, CV 0.2 *p* < 0.001). Notably, within each of the six stimulation groups there was heterogeneity in plasticity outcomes among individual cells (Fig. 4G-H). Importantly, plasticity outcomes in control and mTBI rats were significantly different only when the temporal pattern was highly irregular (*p* = 0.008; *cf.* Fig. 4C, 4F), but not when it was perfectly regular (*p* = 0.76; *cf.* Fig. 4A, 4D) or slightly irregular (*p* = 0.66; *cf.* Fig. 4B, 4E). For highly irregular conditioning control, sham and mTBI groups were significantly different (*p* = 0.004; Fig. 4I). Pairwise comparisons showed mTBI was significantly different from both control (*p* = 0.002) and sham (*p* = 0.002), but control and sham groups did not differ (*p* = 0.49; Fig. 4I). The potentiation produced by highly irregular 1 Hz conditioning in mTBI rats reflects both an increase in the proportion (control = 18%, sham = 6%, mTBI = 47%) and magnitude (post/pre ratio: control = 1.31, sham = 1.13, mTBI = 1.62) of LTP outcomes (Fig. 4I).

**Fig. 4.**
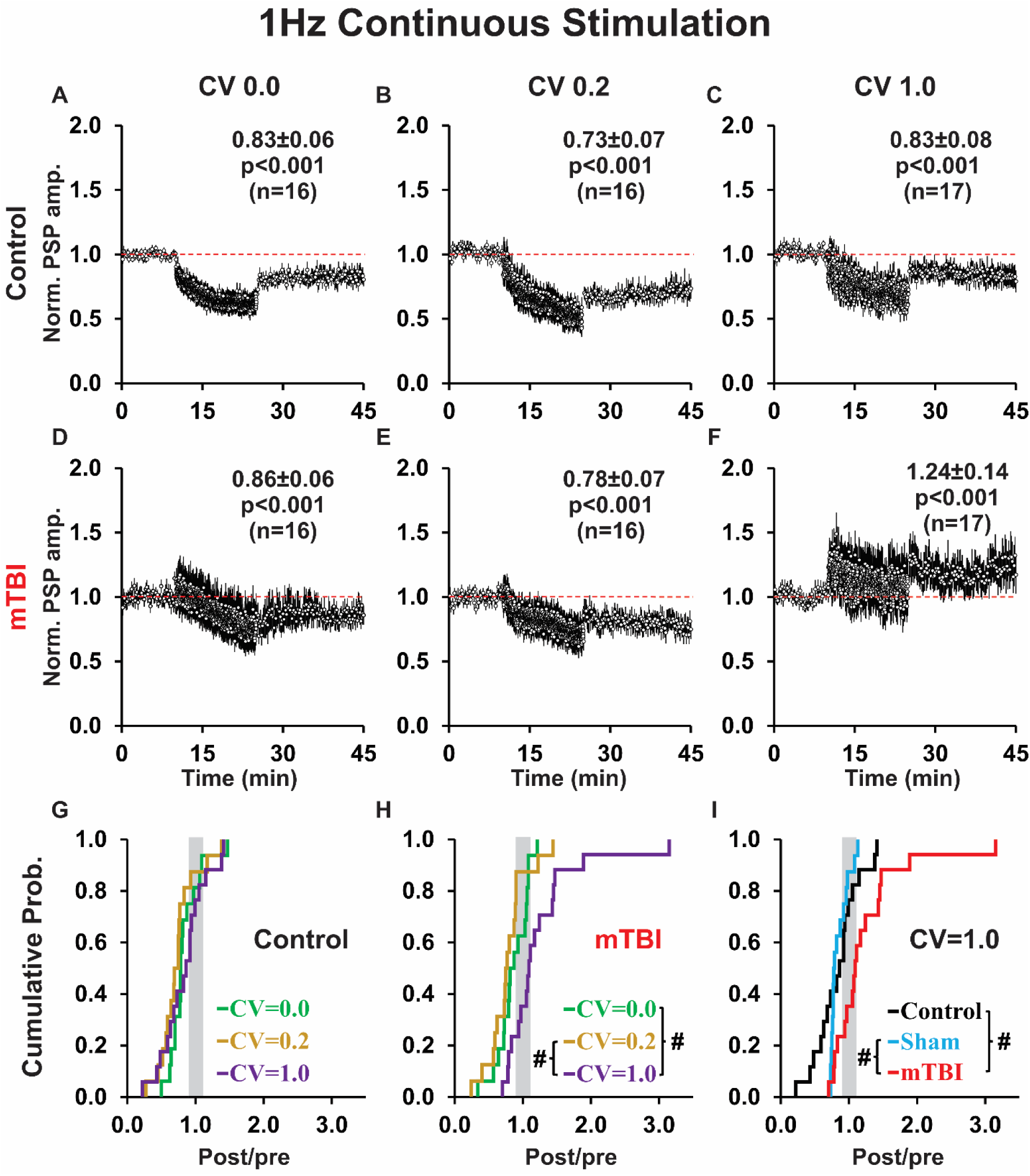
Continuous 1 Hz conditioning induces LTD in every case except one—highly irregular conditioning in mTBI rats results in LTP. (**A-C**) Mean time plots for control groups with: perfectly regular (CV 0.0), slightly irregular (CV 0.2), and highly irregular (CV 1.0) conditioning. Inset text: mean post/pre ratio ± SEM, p-values (*t*-test comparing the last 5 min of pre- vs. post- conditioning), and n = number of cells. Red dashed line = baseline. (**D-F**) Mean time plots for mTBI groups. Note that control and mTBI groups differ for highly irregular (*p* = 0.008 MWU), but not perfectly regular (*p* = 0.76 *t*-test) or slightly irregular (*p* = 0.66 *t*-test) conditioning. (**G-H**) Cumulative probability plots of post/pre ratios in control groups, and mTBI groups. Data steps falling: in the grey region show NC, to the left show LTD, and to the right show LTP. In control groups, temporal pattern had no appreciable effect (*p* = 0.50 ANOVA) on plasticity outcome (curves overlap). However, in mTBI groups, temporal pattern significantly impacted plasticity (*p* = 0.002 Kruskal-Wallis), with pairwise comparisons showing significant potentiation only for highly irregular conditioning: #*p* < 0.017 Dunn’s corrected. (**I**) Cumulative probability plots of post/pre ratio for highly irregular conditioning show a significant effect of treatment group (*p* = 0.004 Kruskal-Wallis, including 16 shams), with subsequent pairwise comparisons revealing a significant shift toward LTP after mTBI: #*p* < 0.017 Dunn’s corrected.

Additionally, highly irregular 1 Hz continuous conditioning differentially affected the evoked PSP shape indices (Fig. S1A) in controls, showing a significant decrease in decay time (DT *p* = 0.03) and half width (HW *p* = 0.03; Fig. S1B), and this effect occurred primarily in the responses that underwent LTD (DT *p* = 0.045, HW *p* = 0.03; Fig. S1C, *cf.* Fig. S1B, S1D). Shape indices in shams showed a similar decrease in DT (*p* = 0.09) and HW (*p* = 0.02; Fig. S1B), which occurred primarily in responses that underwent LTD (DT *p* = 0.03, HW *p* = 0.02; Fig. S1C, *cf.* Fig. S1B, S1D). Interestingly, while the shape indices of the responses in the mTBI group as a whole did not change significantly after conditioning (DT *p* = 0.49, HW *p* = 0.54; Fig. S1B), among just the responses that underwent LTP they significantly increased (DT *p* = 0.01, HW *p* = 0.01; Fig. S1D, *cf.* Fig. S1B-C).

Continuous conditioning at 10 Hz produced a rapid rundown in PSP amplitudes, but was less likely to induce LTD, than 1 Hz continuous conditioning (*cf.* Fig. 5A-E, 4A-E). In particular, for controls, slightly irregular continuous conditioning produced LTD for 1 Hz, but NC for 10 Hz conditioning (*p* = 0.01; *cf*. Fig. 4B, 5B); while after mTBI, perfectly regular conditioning induced LTD for 1 Hz continuous, but LTP for 10 Hz continuous conditioning (*p* = 0.05; *cf*. Fig. 4D, 5D). Temporal pattern had no effect on plasticity outcome in control groups (*p* = 0.58; Fig. 5A-C, 5G). However, for mTBI groups, there was a significant difference in plasticity outcomes (*p* = 0.002; Fig. 5D-F, 5H) between slightly irregular conditioning which showed a shift toward LTD, and perfectly regular conditioning or highly irregular conditioning both of which showed a shift toward LTP (*p* = 0.015; *p* < 0.001; Fig. 5H). Additionally, we found that 10 Hz continuous conditioning (similar to 1 Hz continuous conditioning), produced significantly different plasticity outcomes for control and mTBI groups only when the temporal pattern of conditioning was highly irregular (*p* = 0.003; *cf.* Fig. 5C, 5F), but not perfectly regular (*p* = 0.35; *cf.* Fig. 5A, 5D) or slightly irregular (*p* = 0.10; *cf.* Fig. 5B, 5E). Accordingly, for highly irregular conditioning, we analyzed control, sham and mTBI groups (*p* = 0.003; Fig. 5I). Pairwise comparisons showed a significant potentiation of responses in mTBI relative to control (*p* = 0.001) and sham (*p* = 0.004) groups, but no difference between controls and shams (*p* = 0.30). The potentiated response to highly irregular conditioning (like that observed for the 1 Hz continuous group) reflected a change in both the proportion (control = 0%, sham = 12%, mTBI = 52%) and magnitude (post/pre ratio: control = no LTP responses, sham = 1.24, mTBI = 1.56) of LTP outcomes. However, unlike 1 Hz continuous conditioning, there were no effects of 10 Hz continuous conditioning on the shape indices of PSP responses (data not shown).

**Fig. 5.**
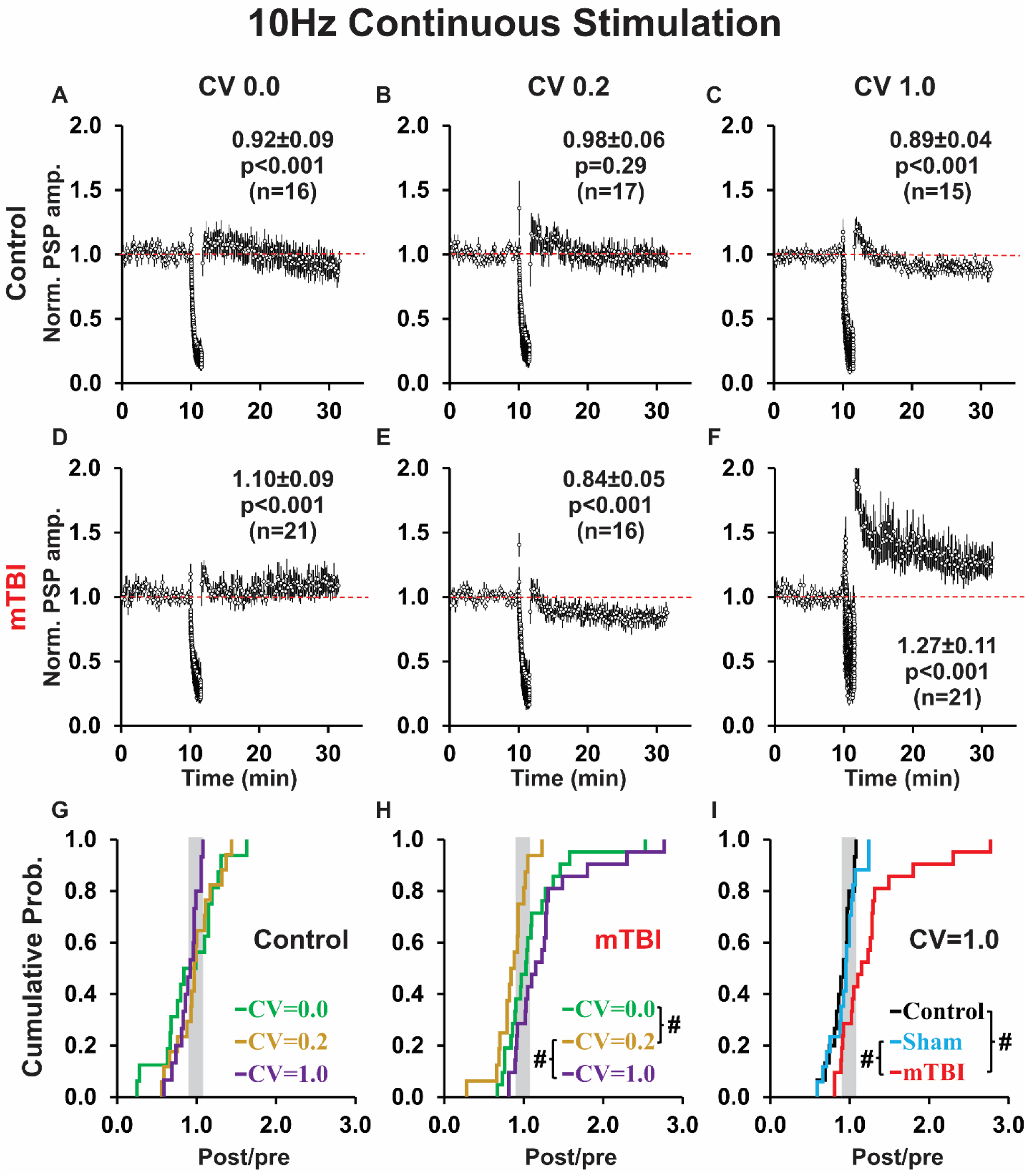
Continuous 10 Hz (compared to 1 Hz) conditioning produces a rapid rundown in PSP amplitudes, is less likely to result in LTD in controls, and more likely to result in LTP after mTBI. (**A-C**) Mean time plots for control groups. (**D-F**) Mean time plots for mTBI groups. Control and mTBI groups differ for highly irregular (*p* = 0.003 MWU), but not perfectly regular (*p* = 0.35 MWU) or slightly irregular (*p* = 0.10 *t*-test) conditioning. (**G-H**) cumulative probability plots of post/pre ratios for: control groups **G**, and mTBI groups **H**. In control groups, temporal pattern has no effect (*p* = 0.58 Kruskal-Wallis) on plasticity outcome (curves overlap). However, in mTBI groups there is a significant effect of temporal pattern (*p* = 0.002 Kruskal-Wallis), with subsequent pairwise comparisons showing significant potentiation for both perfectly regular and highly irregular conditioning: #*p* < 0.017 Dunn’s corrected. (**I**) Cumulative probability plots of post/pre ratio for highly irregular conditioning show a significant effect of treatment group (*p* = 0.003 Kruskal-Wallis, including 17 shams), with subsequent pairwise comparisons revealing a significant shift toward LTP after mTBI: #*p* < 0.017 Dunn’s corrected.

For 10 Hz discontinuous stimulation protocols, the brief conditioning sub-epochs mitigated the rundown of response amplitudes that occurred for 10 Hz continuous conditioning protocols (*cf.* Fig. 6A-F, 5A-F). Moreover, probe stimuli administered at 0.1 Hz during the rest periods (Fig. 6A-F, yellow symbols) showed that PSP amplitudes quickly recovered, in most cases showing a brief period of post-tetanic potentiation, which suggests any synaptic fatigue induced by an individual sub-epoch of higher frequency conditioning was transitory. Interestingly, probe trail response amplitudes were sensitive to the temporal pattern of conditioning in control groups (depressed for CV 0.0 and 1.0, but potentiated for CV 0.2; *cf*. Fig. 6A-C) and mTBI groups (potentiated for CV 0.0, neutral for CV 0.2, and potentiated for CV 1.0; *cf*. Fig. 6D-F). Notably, the continuity of 10 Hz conditioning influenced plasticity outcome and this was also dependent on temporal pattern. In controls, 10 Hz highly irregular conditioning induced a significantly more robust LTD when conditioning was discontinuous rather than continuous (*p* = 0.02; *cf*. Fig. 5C, 6C). In mTBI groups, 10 Hz perfectly regular conditioning induced LTD for discontinuous conditioning, but LTP for continuous conditioning (p = 0.06 - trend; *cf*. Fig. 5D, 6D).

**Fig. 6.**
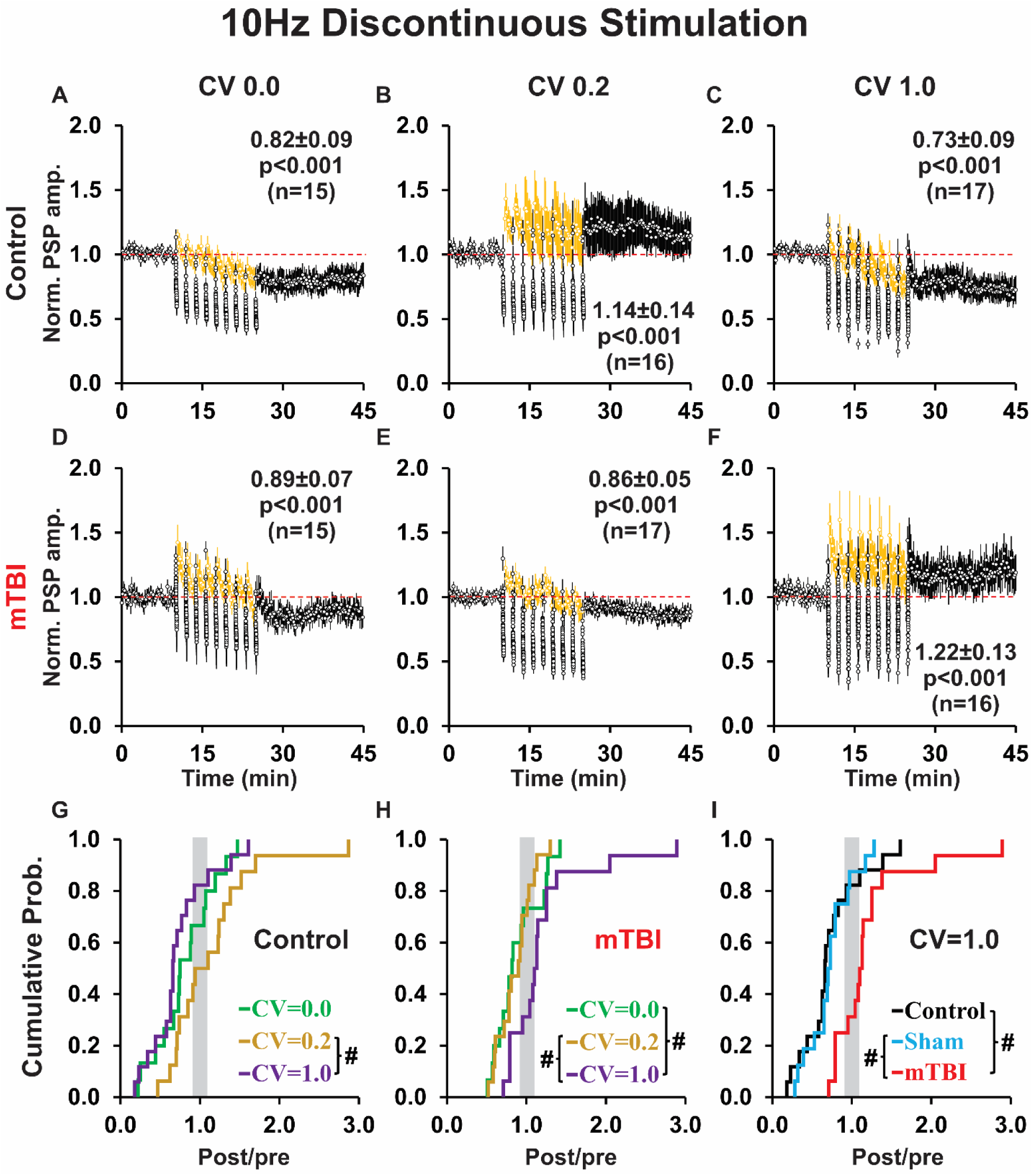
Discontinuous 10 Hz conditioning primarily induced LTD, except for slightly irregular conditioning in controls and highly irregular conditioning in mTBI rats. (**A-C**) Mean time plots for control groups. Note the eight rest periods of 101.25 s (consisting of probe trials at 0.1 Hz, yellow circles), evenly spaced between the nine conditioning sub-epochs (10 s at 10 Hz, black circles). (**D-F**) Mean time plots for mTBI groups. Note that control and mTBI groups differ for highly irregular (*p* < 0.001 MWU), but not perfectly regular (*p* = 0.56 *t*-test) or slightly irregular (*p* = 0.17 MWU) conditioning. (**G-H**) Cumulative probability plots of post/pre ratios for control groups **G**, and mTBI groups **H**. In control groups there is a significant effect of temporal pattern (*p* = 0.04 Kruskal-Wallis), with pairwise comparisons showing a significant difference between slightly irregular conditioning (showing LTP), and highly irregular conditioning (showing LTD): #*p* < 0.017 Dunn’s corrected. In mTBI groups, there is a significant effect of temporal pattern (*p* = 0.02 Kruskal-Wallis), with pairwise comparisons showing a significant shift toward LTP for highly irregular conditioning: #*p* < 0.017 Dunn’s corrected. (**I**) Cumulative probability plots of post/pre ratio for highly irregular conditioning show a significant effect of treatment group (*p* < 0.001 Kruskal-Wallis, including 16 shams), with subsequent pairwise comparisons revealing a significant shift toward LTP after mTBI: #*p* < 0.017 Dunn’s corrected.

For 10 Hz discontinuous (unlike 10 Hz continuous) conditioning there was a significant effect of temporal pattern between control groups (*p* = 0.04; Fig. 6A-C, 6G), as well as between mTBI groups (*p* = 0.02; Fig. 6D-F, 6H). Pairwise comparisons showed a significant effect of temporal pattern in controls, with LTD in response to highly irregular conditioning (Fig. 6C) compared to LTP elicited by slightly irregular conditioning (*p* = 0.006; Fig. 6B, 6G). In mTBI groups, there was a significant difference between highly irregular (Fig. 6F) and both: slightly irregular (*p* = 0.006; Fig. 6E, 6H) and perfectly regular (*p* = 0.012; Figs. 6D, 6H) conditioning. Moreover, control and mTBI groups differed for highly irregular conditioning (*p* < 0.001; *cf.* Fig. 6C, 6F), but not for slightly irregular (*p* = 0.17; *cf*. Fig. 6B, 6E) or perfectly regular (*p* = 0.56; *cf*. Fig. 6A, 6D) conditioning. Accordingly, for highly irregular conditioning, we analyzed control, sham and mTBI groups (*p* < 0.001; Fig. 6I). Pairwise comparisons showed a significant difference between treatment groups—with a robust shift toward LTP in mTBI relative to both control (*p* < 0.001) and sham (*p* <0.001) groups (Fig. 6I). The potentiation reflected an increase in the proportion (control = 12%, sham = 12%, mTBI = 56%), but (unlike 1 Hz and 10 Hz continuous conditioning) not magnitude (post/pre ratio: control = 1.50, sham = 1.23, mTBI = 1.48) of LTP outcomes. There were no effects of 10 Hz discontinuous conditioning on PSP shape indices (data not shown).

For 100 Hz conditioning, we used only a discontinuous protocol which allowed for rapid recovery of PSP amplitudes during rest intervals [with attendant enduring post tetanic potentiation - see yellow probe trial data during conditioning (grey bar) - Fig. 7A-F]. Note that data from the nine conditioning sub-epochs are not shown because most PSPs were embedded in a wave of depolarization and/or had ISIs too short to allow accurate quantification. In control groups, 100 Hz discontinuous perfectly regular and slightly irregular conditioning produced no change in PSP amplitude (*cf.* Fig. 7A, 7B), while highly irregular conditioning induced LTD (Fig. 7C) similar to 1 Hz continuous, 10 Hz continuous and 10 Hz discontinuous conditioning (*cf.* Fig. 7C, 4C, 5C, 6C). In contrast, for mTBI groups, perfectly regular conditioning produced NC (Fig. 7D), slightly irregular conditioning produced LTP [(Fig. 7E) which differed from 1 Hz continuous, 10 Hz continuous, and 10 Hz discontinuous conditioning, all of which exhibited LTD (*cf.* Fig. 7E, 4E, 5E, 6E)], and highly irregular 100 Hz continuous conditioning resulted in NC [Fig. 7F; which differed from 1 Hz continuous, 10 Hz continuous, and 10 Hz discontinuous conditioning—all of which induced LTP (*cf.* Fig. 4F, 5F, 6F)]. Importantly, despite these modest differences, comparisons between control groups (*p* = 0.17), or between mTBI groups (*p* = 0.93) showed no significant effect of temporal pattern on plasticity outcome for the 100 Hz discontinuous conditioning protocol. In addition, there was no significant difference in plasticity outcomes between control and mTBI groups for any temporal pattern (CV 0.0 *p* = 0.40, CV 0.2 *p* = 0.97, CV 1.0 *p* = 0.28).

**Fig. 7.**
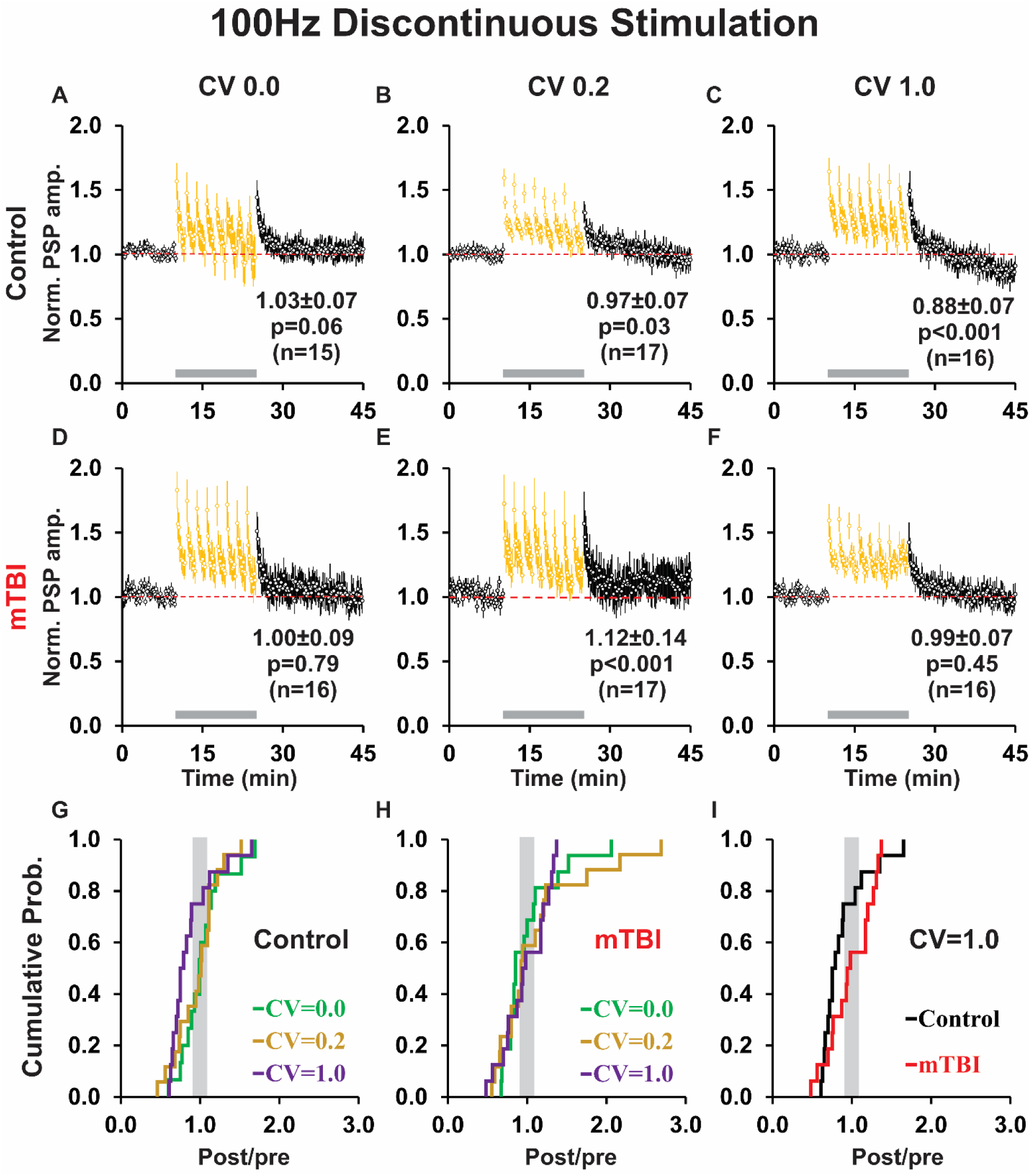
Discontinuous 100 Hz conditioning resulted in little plasticity, except for highly irregular conditioning in controls which produced LTD, and slightly irregular conditioning after mTBI which induced LTP. (**A-C**) Mean time plots for control groups. Grey bars indicate the conditioning period which consisted of nine conditioning sub-epochs (1s at 100 Hz) interspersed between eight evenly spaced rest periods of 131.25 s (with probe trials at 0.1 Hz, yellow symbols). Note that data for conditioning sub-epochs are not shown because the brevity of ISIs precluded measurement of most PSP peak amplitudes. (**D-F**) Mean time plots for mTBI groups. There was no difference in plasticity outcome between control and mTBI groups for any temporal pattern (CV 0.0: *p* = 0.40 MWU, CV 0.2: *p* =0.97 MWU, CV 1.0: *p* = 0.28 *t*-test). (**G-H**) Cumulative probability plots of post/pre ratios for: control groups **G**, and mTBI groups **H**. There was no effect of temporal pattern on plasticity outcome: between control groups (*p* = 0.17 Kruskal-Wallis), or between mTBI groups (*p* = 0.93 Kruskal-Wallis). (**I**) Cumulative probability plots of post/pre ratio for highly irregular conditioning in control and mTBI groups. Note that for 100 Hz discontinuous (unlike 1 Hz continuous, 10 Hz continuous, and 10 Hz discontinuous) highly irregular conditioning, plasticity outcome was not affected by treatment group (control vs. mTBI, *p* = 0.28 *t-test*).

Interestingly, for 100 Hz highly irregular conditioning, the shape indices of PSPs, among the mTBI group showed a significant increase in HW overall (*p* = 0.03; Fig. S1E) that was non-specific for plasticity outcome (LTD *p* = 0.40, LTP *p* = 0.26; *cf.* Fig. S1E-G); while in controls there was no significant change in DT overall (*p* = 0.54; Fig. S1E), but a significant decrease for those cells responding with LTP (*p* = 0.04; Fig. S1G) but not LTD (*p* = 0.39; Fig. S1F). In contrast, for 100 Hz slightly irregular (CV = 0.2) conditioning, the mTBI group showed an increase in RT overall (*p* = 0.002; Fig. S1H) which was consistent across all plasticity outcomes (LTD *p* = 0.035, LTP *p* = 0.006; *cf*. Fig. S1H-J); while in controls RT was unchanged overall (*p* = 0.06) or for any particular plasticity outcome (LTD *p* = 0.18, LTP *p* = 0.47; *cf*. Fig. S1H-J).

### Kinetic properties of conditioning period responses

We analyzed the kinetic profiles of conditioning responses for individual recordings, in the groups that showed a significant difference between control and mTBI, to determine if they were predictive of plasticity outcome. For 1 Hz continuous highly irregular conditioning, we show example time plots of conditioning responses for each treatment group (Fig. 8A-C). Notably, the slope of an exponential best fit of conditioning response amplitudes showed a significant correlation with plasticity outcomes only in control (*p* = 0.02; Fig. 8A, 8D) and sham groups (*p* = 0.048; Fig. 8B, 8D), while the slope of a linear best fit showed a significant correlation with plasticity outcomes only in the mTBI group (*p* = 0.02; Fig. 8C, 8E). The slopes of conditioning response amplitudes did not significantly differ in magnitude for either linear (control = -0.013 ± 0.003, sham = -0.012 ± 0.003, mTBI = -0.007 ± 0.004; *p* = 0.41 ANOVA; data not shown), or exponential best fits (control = -0.019 ± 0.005, sham = -0.018 ± 0.004, mTBI = -0.011 ±0.004; *p* = 0.39 ANOVA; data not shown). However, relative to control and sham groups, the mTBI group exhibited a significant increase in conditioning response amplitude: initial value (overall *p* = 0.002, mTBI vs. control *p* < 0.001, mTBI vs. sham *p* = 0.003), final value (overall *p* = 0.006, mTBI vs. control *p* = 0.001, mTBI vs. sham *p* = 0.007), and average value (overall *p* = 0.01, mTBI vs. control *p* = 0.003, mTBI vs. sham *p* = 0.019—trend; Fig. 8F). Additionally, in the mTBI group, all of these values: average (*p* = 0.004), initial value (*p* = 0.007), and final value (*p* = 0.001) were predictive of plasticity outcome (Fig. 8G-I); while in the control and sham groups, only the average (control *p* = 0.04, sham *p* = 0.03) and final (control *p* = 0.007, sham *p* = 0.009) amplitudes were predictive of plasticity outcome (Fig. 8G, 8I).

**Fig. 8.**
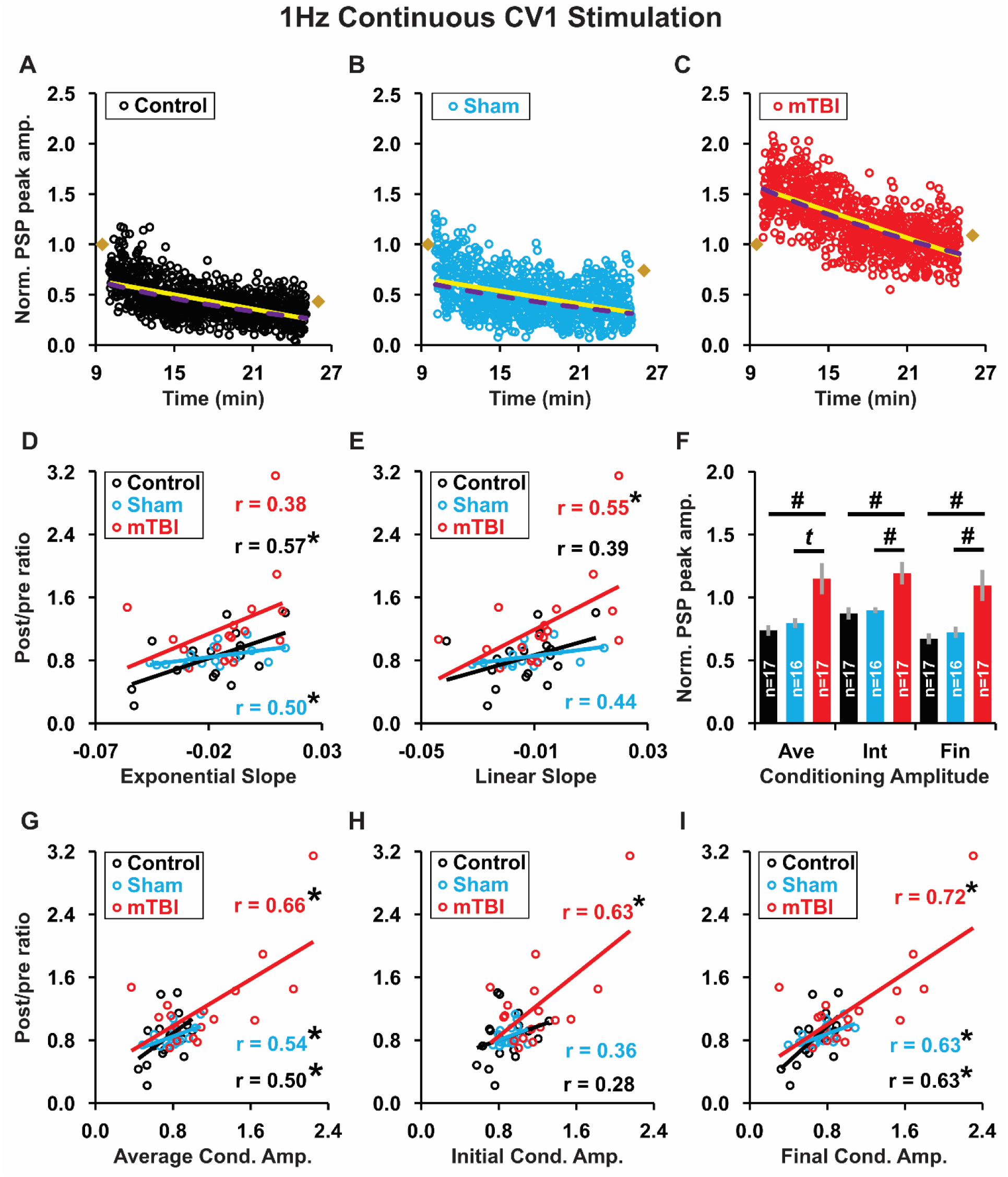
During 1 Hz continuous highly irregular conditioning, the kinetic profile of response amplitudes was predictive of plasticity outcome. (**A-C**) Individual time plots of conditioning period normalized PSP peak amplitudes for representative examples from: control, sham, and mTBI groups. Tan diamonds in each plot show the average of normalized PSP peak amplitudes for the last 5 min of the pre-conditioning (left), and the last 5 min of the post-conditioning (right). For each time plot we calculated the exponential best fit (purple dashed line) and linear best fit (yellow solid line) to obtain the exponential and linear slopes, respectively. (**D-E**) Plots of post/pre ratio (plasticity outcome) verses: exponential slope **D**, and linear slope **E** (control n = 17, sham n = 16, mTBI n = 17; for this and subsequent panels). r = correlation coefficient. Note that the conditioning slope shows a significant correlation for: an exponential fit only in the control and sham groups, and a linear fit only in the mTBI group. \**p* < 0.05 linear regression. (**F**) Normalized PSP peak amplitude during conditioning: Ave (average of all data points), Int (initial 10% of data points averaged), and Fin (final 10% of data points averaged). Values are mean ± SEM. Inset text: n = the number of cells/group. Note that PSP amplitudes are increased in mTBI relative to control and sham groups. #*p* < 0.017 Dunn’s corrected. *t* = trend (*p* = 0.019 Dunn’s corrected). (**G-I**) Plots of post/pre ratio vs. normalized PSP peak amplitude for: average, initial, and final values. Note that while all conditioning amplitudes (average, initial and final) were significantly correlated with plasticity outcome in the mTBI group, only the average and final values were significantly correlated with plasticity outcome in the control and sham groups. \**p* < 0.05 linear regression.

In contrast, for 10 Hz continuous highly irregular conditioning, the kinetics of the conditioning responses (slope Fig. 9A-E; amplitude: Fig. 9G-I) were not predictive of plasticity for any treatment group (control, sham, or mTBI). However, exponential best fits of conditioning amplitude had a more shallow slope in mTBI, than in control or sham groups (overall *p* < 0.001 ANOVA; with mTBI = -0.580 ± 0.096 vs.: control = -1.045 ± 0.090, or sham = -1.125 ± 0.114; *p* = 0.002, *p* < 0.001, Fisher’s LSD; data not shown). Moreover, although the conditioning response amplitude was not significantly different between treatment groups initially (*p* = 0.15; Fig. 9F), both the average (overall *p* = 0.006, mTBI vs. control *p* = 0.009, mTBI vs. sham *p* = 0.002) and final (overall *p* < 0.001, mTBI vs. control *p* = 0.001, mTBI vs. sham *p* < 0.001) amplitudes were significantly higher in the mTBI group than in the control or sham groups (Fig. 9F). Thus, during 10 Hz continuous highly irregular conditioning, response amplitudes decreased less and more slowly, in mTBI than in control and sham groups.

**Fig. 9.**
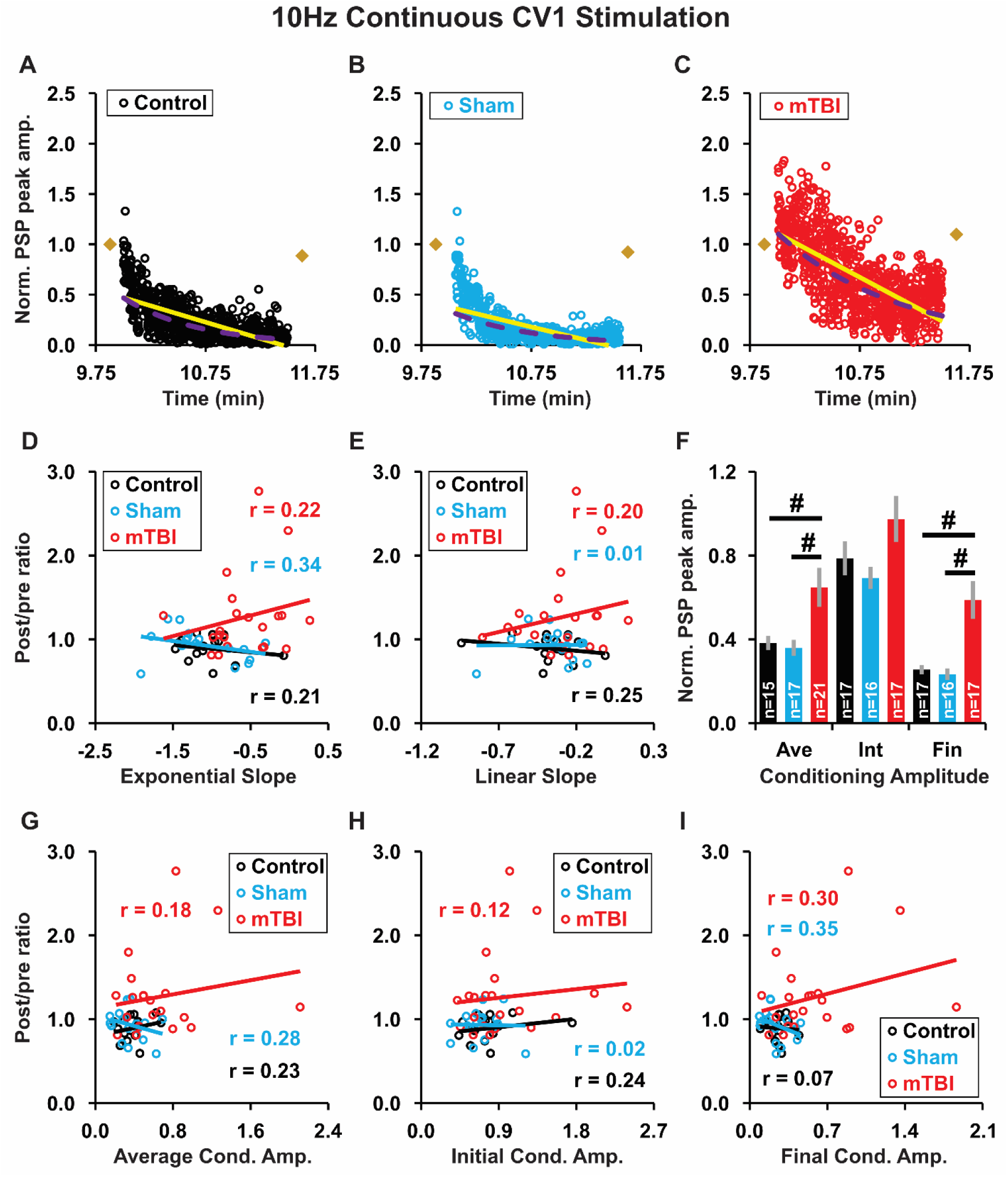
During 10 Hz continuous highly irregular conditioning, the kinetic profile of response amplitudes was not predictive of plasticity outcome. (**A-C**) Individual time plots of conditioning period normalized PSP peak amplitudes for representative examples from: control, sham, and mTBI groups. (**D-E**) Plots of post/pre ratio verses exponential slope **D**, and linear slope **E** (control n = 15, sham n = 17, mTBI n = 21; for this and subsequent panels). r = correlation coefficient. Note that post/pre ratio shows no significant correlation with either exponential or linear slope (controls: *p* = 0.44, *p* = 0.38; shams: *p* = 0.18, *p* = 0.96; mTBI: *p* = 0.34, *p* = 0.37; linear regression). (**F**) Normalized PSP peak amplitude: average (Ave), initial (Int), and final (Fin) values. Note that although the initial amplitude was similar for all treatment groups (*p* = 0.15 Kruskal-Wallis), the average and final amplitudes were significantly increased after mTBI (*p* = 0.006, *p* < 0.001; Kruskal-Wallis). Subsequent pairwise comparisons: #*p* < 0.017, Dunn’s corrected. (**G-I**) Plots of post/pre ratio vs. normalized PSP peak amplitude: average, initial, and final values. Post/pre ratio was not correlated with average, initial or final amplitude for any treatment group (control: *p* = 0.41, 0.38, 0.82; sham: *p* = 0.28, 0.94, 0.18; mTBI: *p* = 0.43, 0.61, 0.19; linear regression).

For 10 Hz discontinuous highly irregular conditioning, the kinetic profile of conditioning response amplitudes measured as the average slope of exponential or linear best fits (Fig. 10A-C), were predictive of plasticity outcome only in control and sham groups—showing significant negative correlations for both linear slope (control *p* < 0.001, sham *p* = 0.02), and exponential slope (control *p* < 0.001, sham *p* = 0.04; Fig. 10D-E). This result differs from both 1 Hz continuous highly irregular conditioning (significant positive correlation, *cf.* Fig. 8D-E), and 10 Hz continuous highly irregular conditioning (no correlation, *cf.* Fig. 9D-E). There was no significant difference in the magnitude of conditioning response slopes between treatment groups, for either: linear (control -0.560 ± 0.376, sham -0.908 ± 0.233, mTBI -1.157 ± 0.155; *p* = 0.19 Kruskal-Wallis; data not shown) or exponential best fits (control -0.899 ± 0.671, sham -1.555 ± 0.297, mTBI -1.959 ± 0.388; *p* = 0.24 Kruskal-Wallis; data not shown). Additionally, there was no significant difference in the magnitude of conditioning response amplitude: average (*p* = 0.75), initial (*p* = 0.95), or final values (*p* = 0.41; Fig. 10F). However, in the mTBI group, conditioning response amplitude: average (*p* = 0.02), initial value (*p* = 0.03) and final value (*p* = 0.02) were all predictive of plasticity outcome (Fig. 10G-I). In contrast, for the control and sham groups, conditioning response amplitude was not predictive of plasticity outcome for: average (control *p* = 0.71, sham *p* = 0.35; Fig. 10G), initial (control *p* = 0.17, sham *p* = 0.24; Fig. 10H), or final values (control *p* = 0.69, sham *p* = 0.44; Fig. 10I).

**Fig. 10.**
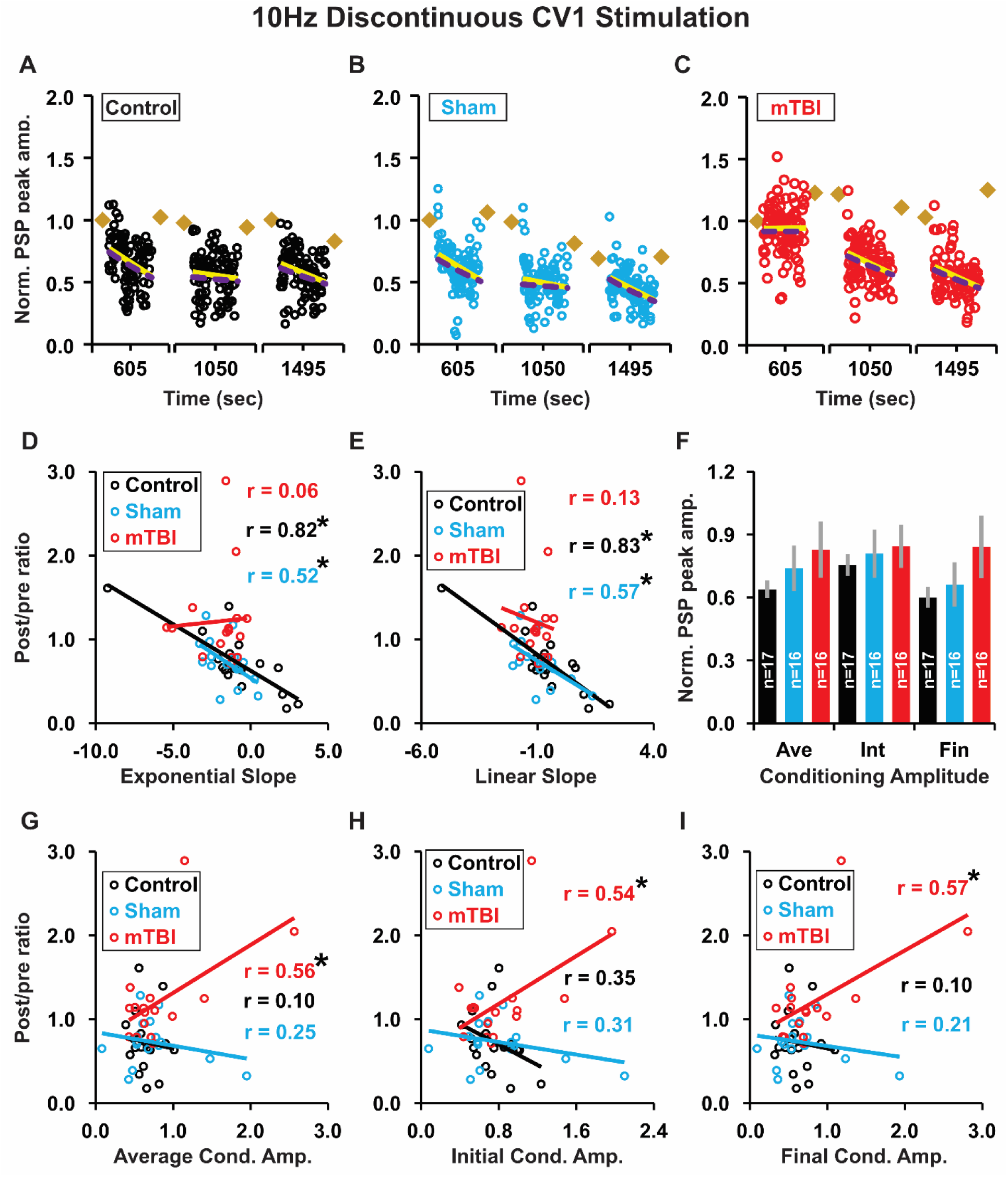
During 10 Hz discontinuous highly irregular conditioning, the kinetic profile of PSP responses had: slopes (exponential and linear) that were significantly correlated with plasticity outcome in control and sham, but not mTBI groups; and amplitudes (average, initial and final) that were significantly correlated with plasticity outcome in mTBI groups, but not control or sham groups. (**A-C**) Individual time plots of conditioning period normalized PSP peak amplitudes for representative examples of: control, sham, and mTBI groups. Note that only the first (left), fifth (center) and ninth (right) conditioning sub-epochs are shown. Tan diamonds represent the average of normalized PSP peak amplitudes for (from left to right): the last 5 min of pre-conditioning and the first rest interval (left), the fourth and fifth rest intervals (center), and the eighth rest interval and the last 5 min of post-conditioning (right). For each conditioning sub-epoch we calculated the exponential best fit (purple dashed line) and linear best fit (yellow solid line) to obtain the exponential and linear slopes. (**D-E**) Plots of post/pre ratio vs. conditioning slope (average of all 9 conditioning sub-epochs) for exponential **D**, and linear **E** best fits, respectively (control n = 17, sham n = 16, mTBI n = 16; for this and subsequent panels). r = correlation coefficient. Note that the conditioning slopes (exponential and linear) show a significant negative correlation with plasticity outcome in control (*p* < 0.001, *p* <0.001) and sham (*p* = 0.04, *p* = 0.02), but not mTBI (*p* = 0.84, *p* = 0.64) groups. \**p* < 0.05, linear regression. (**F**) Normalized PSP peak amplitude: average (Ave), initial (Int), and final (Fin) values. Treatment group (control, sham, mTBI) had no effect on average (*p* = 0.75 Kruskal-Wallis), initial (*p* = 0.95 Kruskal-Wallis) or final amplitude (*p* = 0.41 Kruskal-Wallis). (**G-I**) Plots of post/pre ratio vs. normalized PSP peak amplitude: average, initial, and final values. Conditioning amplitudes (average, initial and final) showed a significant positive correlation with plasticity outcome in the mTBI (*p* = 0.03, 0.04, 0.03), but not the control (*p* = 0.71, 0.17, 0.69) or sham group (*p* = 0.35, 0.24, 0.44; linear regression). \**p* < 0.05 linear regression.

### Data from our WC recordings appear broadly representative

In order to assess whether our WC results were reflective of the larger population of synapses in L2/3, we also made simultaneous FP recordings. Representative examples of individual recordings and time plots for paired WC and FP recordings are shown in Supplementary Figure 2 for: 1 Hz continuous (Fig. S2A-B), 10 Hz continuous (Fig. S2C-D), and 10 Hz discontinuous (Fig. S2E-F) conditioning. Notably, plasticity was highly correlated between the simultaneous WC and FP recordings regardless of treatment group (control *p* < 0.001, sham *p* < 0.001, mTBI *p* < 0.001; Fig. S2G) or temporal pattern (CV 0.0 *p* < 0.001, CV 0.2 *p* < 0.001, CV 1.0 *p* < 0.001; Fig. S2H).

## DISCUSSION

We found that mTBI dramatically affected the dependency of plasticity on the temporal pattern of conditioning. At all frequencies studied, with highly irregular temporal patterns of synaptic conditioning, where neurons from controls underwent LTD (Fig. 4C, 5C, 6C, 7C), those from mTBI animals underwent LTP (for 1 Hz or 10 Hz; Fig. 4F, 5F, 6F) or NC (for 100 Hz; Fig. 7F). Moreover, with slightly irregular or perfectly regular temporal patterns of conditioning, neurons from mTBI animals underwent different plasticity outcomes than those from controls (LTP vs. NC for 10 Hz continuous conditioning, CV 0, *cf.* Fig. 5D, 5A; LTD vs. NC for 10 Hz continuous conditioning, CV 0.2, *cf.* Fig. 5E, 5B; and LTD vs. LTP for 10 Hz discontinuous conditioning, CV 0.2, *cf.* Fig. 6E, 6B). Finally, for slightly or highly irregular 100 Hz discontinuous conditioning, neurons from mTBI rats displayed different plasticity outcomes from controls (LTP vs. NC for CV 0.2, *cf.* Fig. 7E, 7B; and NC vs. LTD for CV 1.0, *cf.* Fig. 7D, 7A).

To our knowledge previous studies have not examined the effect of temporal pattern on the induction of LTP or LTD following TBI, but they have shown that temporal pattern effects induction of synaptic plasticity in the normal brain. Studies using 2 Hz Markovian stimulation with different second order statistics reported strong negative correlations produced LTD, while more positive correlations produced LTP.^110–112^ Additionally, studies using a spike timing dependent plasticity protocol evoked by paired pre-post synaptic stimulation at 0.5–3 Hz, observed LTD if ISIs were regular or had a Gaussian distribution, but no plasticity (NC) if ISIs had a Poisson distribution.^113,114^ A direct comparison with these studies is difficult because we used a different approach in generating our stimulation protocols, but one study^115^ used a similar approach— varying the CV for ISIs—and like them we found that 1 Hz perfectly regular conditioning produced LTD, but unlike them we found that 1 Hz highly irregular conditioning often did too. This discrepancy in plasticity outcomes for highly irregular conditioning may reflect subtle but important differences in the temporal pattern of ISIs in the two studies; or could reflect differences in: stimulus pulse duration, species, or degree of brain maturation including the development of synaptic inhibition.

### Analyses of PSP shape index and conditioning response kinetics suggest underlying plasticity mechanisms in mTBI

Our WC patch clamp recordings of compound PSPs from L2/3 pyramidal neurons evoked by stimulation of L4 probably included a mixture of excitatory and inhibitory components to L2/3 pyramidal neurons.^116–120^ Therefore changes in evoked PSP amplitude after conditioning could reflect an alteration in any of these components. For example, LTP of the PSP could be due to an increase in the strength of excitatory transmission, a decreased strength of inhibitory transmission, or both.

We chose not to pharmacologically isolate either component as we aimed to have the experiments directly address the type of neurostimulation that would be used in a clinical situation. However, we did apply an analysis of PSP shape indices^115,121–124^ (half width, rise time and decay time) to our WC recordings (Fig. S1). These results were generally consistent with LTP of the PSP being due to an increase in excitatory transmission strength and LTD being due to a decrease in excitatory transmission strength. Moreover, the FP results (Fig. S2G-H) that are generally thought to reflect excitatory synaptic transmission^125,126^ were consistent with this result.

Our results show that the frequency, continuity and temporal pattern of conditioning stimulation interact to modify synaptic plasticity, but that interaction of these factors produces different plasticity outcomes in control (naïve and sham) rats verses those with mTBI. Application of these results can also be informed by our findings on the utility of the kinetic profile of synaptic responses during the conditioning (Fig. 8-10) for predicting a desired plasticity outcome. Our utilization of probe trials (Fig. 6-7) to assess synaptic strength throughout the conditioning period could be useful to adapt a treatment protocol in real time to achieve the desired plasticity outcome.^127–132^

The increased intracellular calcium concentration and input resistance we observed following mTBI (Fig. 2) are indicative of increased neuronal excitability.^133,134^ Evidence suggests that differences in the magnitude, duration, rate of rise, source and spatial distribution of intracellular calcium signals are important for triggering LTD vs. LTP.^129,135–141^ These distinct calcium signals may act differentially on downstream signaling cascades,^142–146^ to selectively alter postsynaptic AMPA receptors^147–152^ and translation/transcription factors.^153–158^ Moreover, intracellular calcium levels can increase over larger spatial and temporal scales, for example via entry through voltage-gated calcium channels and/or release from intracellular stores,^159–163^ which can affect processes such as gene expression and protein synthesis that contribute to more persistent synaptic plasticity.^164–167^

### Implications of our results for potential therapeutic interventions in the treatment of mTBI

One potential application of our findings is to inform design of stimulation parameters for clinical interventions such that different patterns of synaptic conditioning characterized by specific combinations of temporal pattern, frequency, and continuity, could be deployed to modulate plasticity outcomes, strengthening or weakening affected neural circuits, to promote recovery of cortical function after mTBI. Some studies^168–172^ in humans with implanted stimulating electrodes have shown dramatic results, although the relationship of various stimulation parameters to treatment efficacy are poorly understood.^173^ Moreover, advances in technology such as the use of low intensity MRI guided transcranial focused ultrasound for neuromodulation^174–176^ provide an opportunity to consider such potential therapeutic stimulation protocols without invasive brain surgery.

## TRANSPARENCY, RIGOR AND REPRODUCIBILITY SUMMARY

The study design and analytical plans were not formally pre-registered, but they were pre-specified. The planned sample size for the righting reflex experiment was 12-15 rats/treatment group (control, sham, mTBI) based on data from similar studies.^177,178^ For the electrophysiology experiments the planned sample size was 15-20 cells for each of the 24 experimental groups (4 stimulation protocols—1 Hz continuous, 10 Hz continuous, 10 Hz discontinuous, 100 Hz discontinuous; 3 temporal patterns—perfectly regular, slightly irregular, highly irregular; and 2 treatment groups—control, mTBI) based on data from similar studies.^115,138,179^ Given that the typical yield of successful WC recordings in these studies was about 1.5 cells/animal, we estimated that 320 rats would be required for the electrophysiology experiments. Additionally, when control and mTBI results for four of the stimulation groups appeared divergent, we amended our experimental design to include 55 sham operated controls.

The actual number of rats used for the righting reflex experiments was 45 (including 1 sham excluded for a disruption during testing); while that for the electrophysiology experiments was 389 rats (including: 1 sham and 5 mTBI rats excluded for surgical complications; and 67 rats excluded from WC experiments because recordings were incomplete, but retained for calcium experiments because pre-conditioning was successful). From these animals we obtained 446 successful WC recordings and 222 successful calcium imaging experiments. This corresponded to 15-21 cells/experimental group for WC experiments, and 37-96 cells/treatment group for calcium imaging experiments. Statistical significance was determined using SigmaPlot (Systat Software) or an online service (statskingdom.com/kruskal-wallis-calculator.html). Normality was assessed with a Shapiro-Wilk test. For normally distributed data we used a *t-test*, or a one-way ANOVA followed by Fisher’s LSD method for multiple comparisons. For non-normally distributed data we used a MWU test, or a Kruskal-Wallis test followed by Dunn’s method for multiple comparisons with Sidak correction (Dunn’s corrected). Differences were considered statistically significant at *p* < 0.05, except for Dunn’s corrected which was considered significant at p < 0.017.

At 8-9 weeks of age, rats were randomly assigned to treatment groups by an investigator who also performed the surgeries (mTBI or sham). Two to three weeks later a different set of investigators blinded to treatment group performed the righting reflex, electrophysiology, and calcium imaging experiments. All data collection and analysis of individual experiments (including criteria-based exclusions—see methods) were performed by the investigators blinded to treatment group.

Data are available from the corresponding author upon request. The authors have agreed to publish this paper under a Creative Commons Open Access license, and upon publication it will be freely available at: liebertpub.com/loi/neu.

## ACKNOWLEDGEMENTS

We would like to thank Susanna Kiss for administrative and technical support with: blinding procedures, sham and mTBI surgeries, righting reflex experiments, and histological processing. Additionally, we would like to express our gratitude to: members of Dr. Claudia Robertson’s laboratory for instruction and technical assistance with mTBI and sham procedures, and Dr. P. Read Montague for helpful discussions regarding the development of temporal pattern distributions and conceptualization of the experiments.

## AUTHORS’ CONTRIBUTIONS

Conceptualization: QSF, DK, GVDP, PRB, MJF; Methodology: QSF, DK, GVDP, PRB, MJF; Validation: QSF, DK; Formal analysis: QSF, DK, GVDP, CAW; Investigation: QSF, DK, GVDP, CAW; Software: QSF, CAW, PRB; Data curation: QSF, CAW, PRB; Writing – original draft: QSF, MJF; Writing – review and editing: QSF, DK, GVDP, CAW, PRB, MJF; Visualization: QSF, DK, MJF; Supervision: QSF, MJF; Project administration: QSF, MJF; Resources: MJF; Funding Acquisition: MJF. All authors contributed to the manuscript and approved the submitted version.

## SUPPLEMENTARY MATERIALS

Supplementary Figure S1

Supplementary Figure S2

Detailed Methods

