## Supplementary Materials for "SYNAPTIC PLASTICITY IN THE INJURED BRAIN DEPENDS ON THE TEMPORAL PATTERN OF STIMULATION"

**Supplementary Fig. S1. Significant changes in shape indices were induced by only a few stimulation protocols, and were dependent on treatment group, temporal pattern, and plasticity outcome.** (**A**) Example PSP waveforms averaged over the last 5 min of pre- (grey) or post- (orange) conditioning. Red dotted line = baseline. Peak/solid arrows = PSP peak amplitude; RT = rise time (10-90% of peak amplitude), DT = decay time (90-10% of peak amplitude), arrowheads indicate 10-90% bounds of RT or DT; HW / open arrows = half width (at half height). (**B-D**) PSP shape indices before (pre-conditioning, solid bars) and after (post-conditioning, open bars) 1 Hz continuous highly irregular conditioning for: all cells, only cells showing LTD, and only cells showing LTP. n = number of cells/treatment group (control black, sham blue, mTBI red)—for this and subsequent panels. *$p < 0.05$ paired *t*-test (for this and subsequent panels). (**E-G**) PSP shape indices before and after 100 Hz discontinuous highly irregular conditioning for: all cells, only cells showing LTD, and only cells showing LTP. (**H-J**) PSP shape indices before and after 100 Hz discontinuous slightly irregular conditioning for: all cells, only cells showing LTD, and only cells showing LTP. **$p < 0.01$ paired *t*-test.

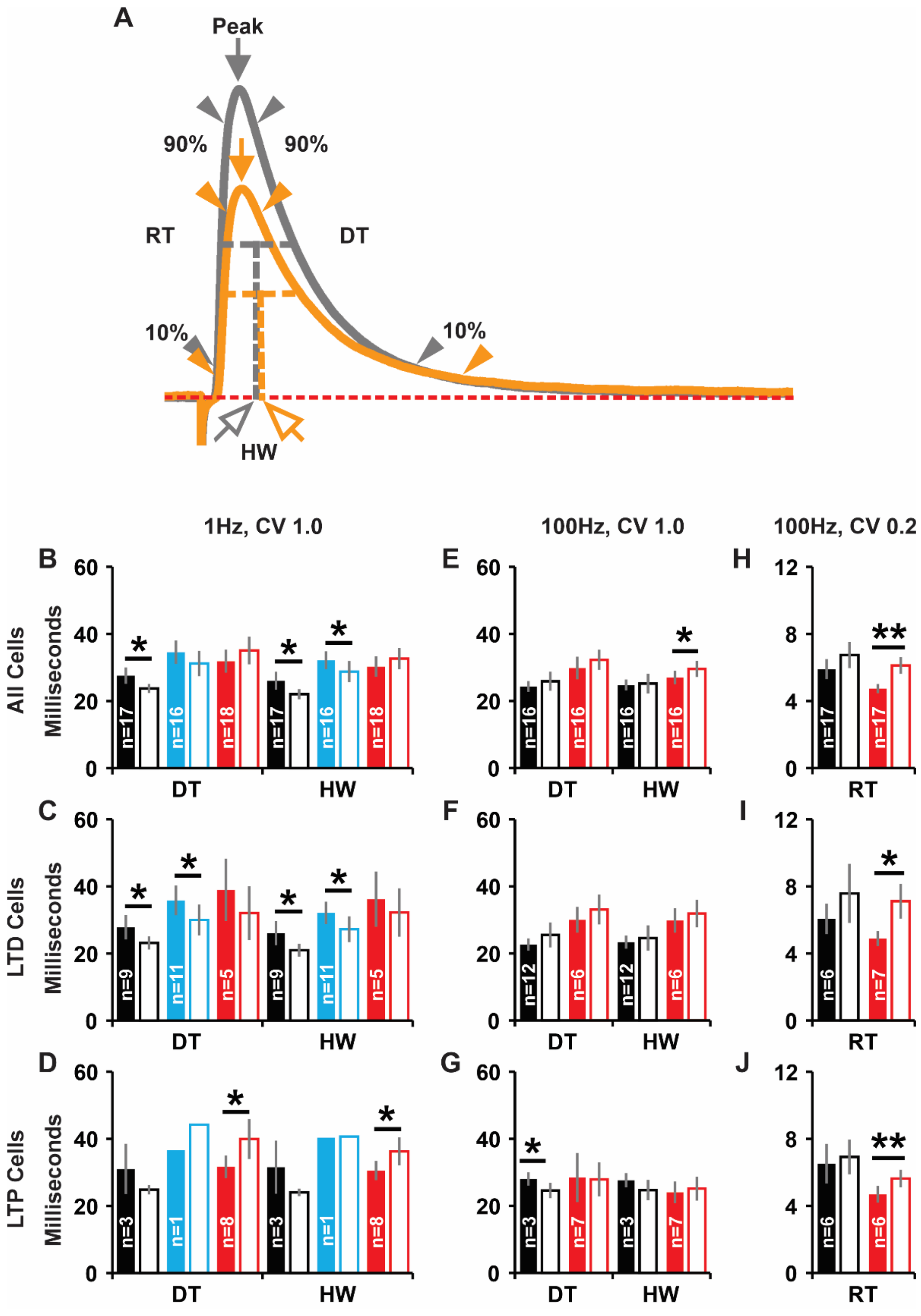


**Supplementary Fig. S1**

**Supplementary Fig. S2. Individual whole-cell and field potential recordings made in parallel show a significant correlation in plasticity outcomes.** (**A-B**) Individual time plots of WC and FP responses to 1 Hz continuous conditioning (black open circles = pre-/post-conditioning period responses, black filled circles = conditioning period responses). Inset text: mean post/pre ratio ± SEM, with the corresponding plasticity outcome (LTD, NC or LTP). Inset figure: averaged response waveforms for the last 5 min of pre- (grey) and post- (orange) conditioning (scale bars: WC = 5 mV, 50 ms; FP = 50 μV, 10 ms). WC and FP time plots shown on the same row were recorded in parallel from the same slice. (**C-D**) Individual time plots of WC and FP responses to 10 Hz continuous conditioning. (**E-F**) Individual time plots of WC and FP responses to 10 Hz discontinuous conditioning. Yellow symbols during conditioning represent probe trials (at 0.1 Hz) between stimulus epochs. All examples **A-F** depict control data. (**G-H**) WC and FP recordings made in parallel from the same slice have post/pre ratios which are significantly correlated regardless of treatment group: control (n = 50), sham (n = 26), or mTBI (n = 69); or temporal pattern CV 0.0 (n = 25), CV 0.2 (n=43), or CV 1.0 (n=77). ***$p < 0.001$ linear regression. Solid lines represent a linear best fit, r = correlation coefficient.

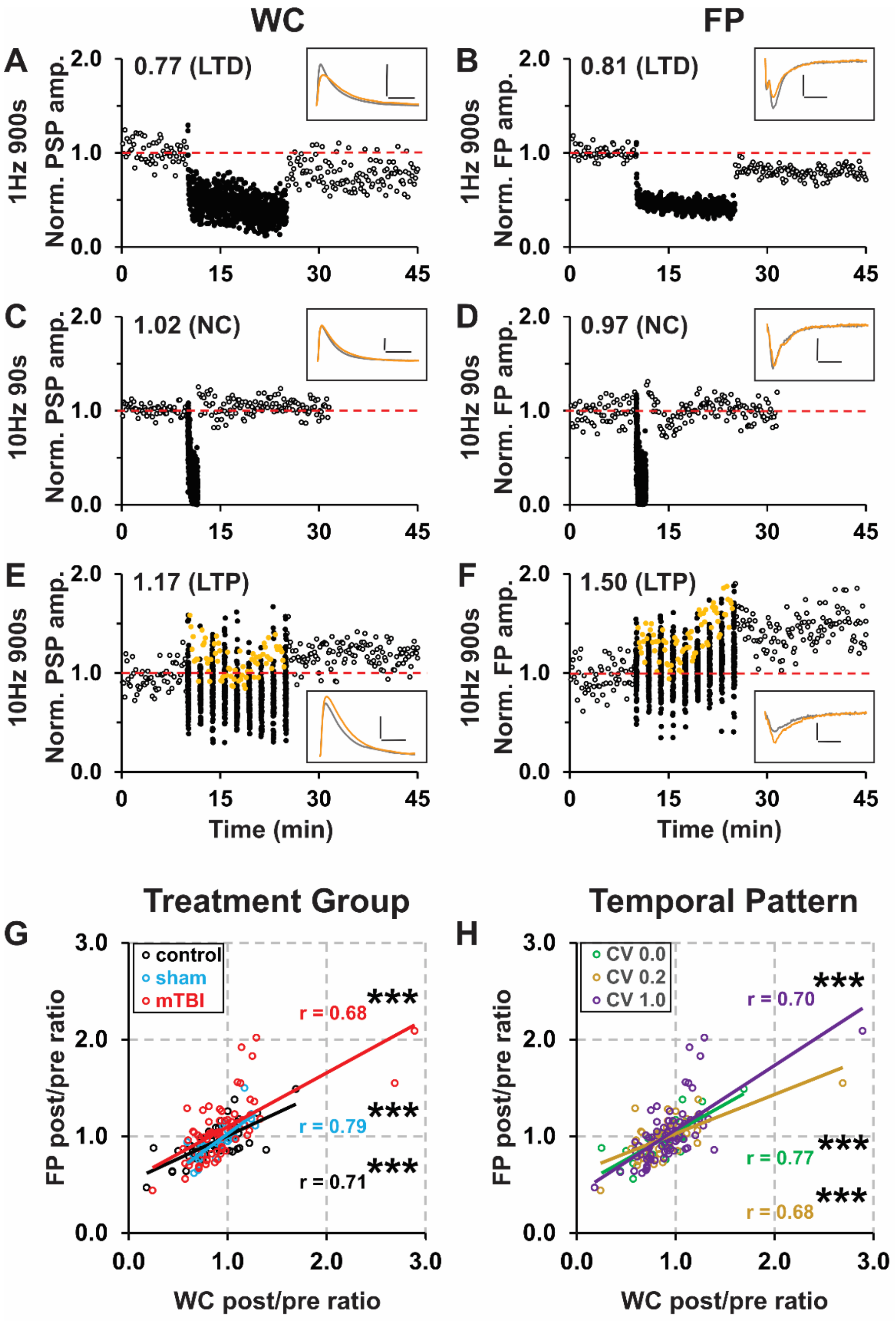


Supplementary Fig. S2

### DETAILED METHODS

#### Animal use and experimental design

All procedures were approved by the Institutional Animal Care and Use Committees of Virginia Tech or Baylor College of Medicine and followed the National Research Council's *Guide for the Care and Use of Laboratory Animals*. Male Long-Evans rats (HsdBlu:LE, Inotiv) were kept on a 12h light / 12h dark schedule, with food and water *ad libitum*. At 8-9 weeks of age, rats were single-housed and assigned to one of three treatment groups: control (naïve), sham (sham operated), or mTBI (controlled cortical impact, CCI). Two to three weeks later these animals were used in: behavioral, electrophysiological, and/or calcium imaging experiments. Data collection, analysis and exclusion of individual experiments were performed by investigators blinded to treatment group.

#### Mild traumatic brain injury model

At 8-9 weeks of age the mTBI group received a CCI using a well-established model.[1–3] Four hours before surgery rats were started on an antibiotic regimen (Baytril 5 mg/kg, IM, *q.d.* x 3). Surgery was performed under isoflurane anesthesia (1–4%, in $O_2$). Rats were intubated, ventilated (TOPO, Kent Scientific), and end tidal $CO_2$ was monitored (Capnograph 340, Harvard Apparatus). Animals were placed on a homeostatically controlled heating plate (TCAT-2DF, Physitemp) and mounted in a stereotaxic frame (Model 940, Kopf). The scalp was shaved, cleaned with alcohol and betadine, and incised to expose the skull. A 7 mm diameter craniotomy was made over the right parietal cortex immediately posterior to bregma and lateral to midline. A direct lateral CCI (2.5 mm depth, 3.0 m/s velocity, 100 ms duration, angled at 45° from vertical) was delivered through intact meninges using a Benchmark stereotaxic impactor (Leica) with a modified 6 mm diameter spherical head. After the CCI, we administered a long-lasting analgesic (buprenorphine SR 1.2 mg/kg, SC; ZooPharm), the craniotomy and incision were repaired, and a topical antibiotic (Neosporin, Johnson & Johnson) was applied. Anesthesia was discontinued, the rat removed from the stereotaxic frame, extubated, and monitored until fully recovered. Shams received identical treatment except for the CCI.

**Righting reflex**

At 10-12 weeks of age (2-3 weeks after treatment: control, sham, or mTBI) the first cohort of 45 rats was examined to assess righting latency. Individual rats were placed in a transparent induction chamber (with 1L/min $O_2$). Anesthesia was induced by the addition of isoflurane (4% for 120 s). The rat was rapidly removed from the chamber and placed in a supine position. The time between the cessation of anesthesia and the recovery of the righting reflex (righting latency, defined as the time required for the rat to right itself to an upright posture standing on all four paws) was measured.

**Brain slice preparation**

At 10-12 weeks of age (2-3 weeks after treatment: control, sham, or mTBI) a second cohort of 389 rats was anesthetized with ketamine / xylazine (75 and 10 mg/kg, IM) then decapitated. The brain was rapidly removed and cooled for ~2 min in ice-cold cutting solution containing (in mM): 124 NaCl, 2 KCl, 3.5 $MgCl_2$, 0.2 $CaCl_2$, 1.25 $KH_2PO_4$, 26 $NaHCO_3$, 11 dextrose, with pH 7.4, and saturated with carbogen (95% $O_2$+5% $CO_2$). Coronal slices (300 μm) were prepared from the right visual cortex (ipsilateral to the mTBI) using a vibratome (VT1000P, Leica) and placed in artificial cerebral spinal fluid (ACSF—containing in mM: 124 NaCl, 2 KCl, 2 $MgCl_2$, 2 $CaCl_2$, 1.25

$KH2PO_4$, 26 $NaHCO_3$, 11 dextrose, with pH 7.4, and saturated with carbogen). Slices were incubated in ACSF at 33 ± 1ºC for 20 min and then at room temperature for > 40 min prior to recording.

### ***In vitro*** **electrophysiology**

For recording, slices were transferred to a submersion chamber perfused with 3 ml/min of ACSF at 33±1ºC. L2/3 pyramidal neurons were visualized using a Zeiss Axio Examiner D1 microscope equipped with a W Plan Apochromat 40x water immersion lens and configured for DGC microscopy. A bipolar stimulating electrode (FHC #30213) was placed in L4. Glass micropipettes (Garner KG-33, OD 1.5 mm, ID 1.12 mm; A-M Systems) were pulled on a vertical puller (PC-10, Narishige) to make FP pipettes (1 ± 0.5 MΩ), or WC patch pipettes (4 ± 1 MΩ). FP pipettes were filled with ACSF and placed superficially in L2/3. WC patch pipettes were filled with pipette solution (containing in mM: 115 K-gluconate, 20 KCl, 10 Hepes, 4 NaCl, 4 Mg-ATP, 0.3 Na-GTP, and 4 phosphocreatine-Na; pH 7.4; 280-290 mOsm) and placed in L2/3 (~100 µm deep to the FP pipette). Biocytin (0.5%; Sigma-Aldrich) was added to the WC patch pipette solution to allow post-hoc identification of patched cells (only pyramidal cells located in L2/3 were analyzed).

All recordings were made using a MultiClamp 700B amplifier, digitized at 20 kHz with a Digidata 1440A, and collected and analyzed with pCLAMP 10 software (Axon Instruments, Molecular Devices). Recordings of evoked PSP and FP amplitudes were measured as the difference between baseline and peak. Recordings with unstable pre-conditioning response amplitudes (linear best fit of PSP, or FP, peak amplitudes with an $r^2 > 0.2$) were discarded. WC recordings with membrane potentials more positive than -60 mV, high access resistance (> 40 MΩ, or > 20% of the membrane resistance for that cell), or a > 20% decrease in input resistance were discarded. The intrinsic firing properties of patched cells were tested by injecting a 100 ms depolarizing current pulse and cells that did not exhibit regular spiking typical of excitatory cells[4,5] were discarded. Only one recording was made from each slice, so that a single stimulation protocol was applied in each case.

**Stimulation protocols**

Afferent stimulation consisted of constant current square wave pulses (100 μs in duration, and 30-500 μA in amplitude; set to evoke ~40% of the maximal PSP response) output by a stimulus isolation unit (SIU; A365, World Precision Instruments). Stimulation was optimized to produce PSP responses, but in some cases (~30%) also evoked FP responses which were recorded simultaneously. Pre-conditioning baseline recordings of evoked PSP (and FP) peak amplitudes were made at 0.1 Hz for 10 min. This was followed by a 90 or 900 s period of synaptic conditioning, consisting of 900 pulses at 1 Hz, 10 Hz, or 100 Hz. Conditioning stimulation was either continuous (1 Hz for 900 s, or 10 Hz for 90 s; Fig. 1A-B) or discontinuous (9 epochs of 100 pulses each at 10 Hz or 100 Hz, separated by 8 equal rest intervals consisting of probe trials at 0.1 Hz, with a total duration of 900 s; Fig. 1C). We included the discontinuous paradigms since continuous stimulation at higher frequencies (10 Hz or more) can induce presynaptic fatigue that might interact with long-term synaptic plasticity.[6,7] Post-conditioning evoked PSP (and FP) peak amplitudes were evaluated at 0.1 Hz for 20 min. Conditioning stimulation was delivered with one of three temporal patterns: perfectly regular (all ISIs equal, with a coefficient of variation = 0, CV 0.0; Fig. 1D-E), slightly irregular (Gaussian distribution of ISIs, CV 0.2; Fig. 1F-G), or highly irregular (Poisson distribution of ISIs, CV 1.0; Fig. 1H-I) each with the same mean frequency. Conditioning stimulation was generated using MatLab (MathWorks), transformed with a digital analog converter (BNC 2090A, National Instruments), triggered with a pulse stimulator (Master-9, A.M.P.I.), and output via the SIU, giving us precise control of frequency, continuity and temporal pattern.

**Temporal patterns of conditioning stimulation**

Conditioning stimulation temporal patterns were generated by creating ISIs according to the family of distributions given by:

$$P(t) \propto e^{-t^2/2\,\eta^2}\; e^{-t\,/\tau_d}\;\; \theta(t > \tau_R) \qquad (1)$$

where $\tau_R$ is the refractory period, and $\eta$ (>0) and $\tau_d$ are temporal parameters that govern the single peak of the distribution. For a perfectly regular distribution, we had $-\tau_d/\eta >> 1$, and for a highly irregular distribution $\eta/\tau_d >> 1$. We solved for the parameters $\eta$ and $\tau_d$, to yield a desired mean, m, and standard deviation (std) or coefficient of variation (CV):

$$\frac{\mathrm{std}^2}{\mathrm{m}^2} = \frac{Q(w)^2 + 2wQ(w) - 2}{2(1 - wQ(w))^2}, \quad (2)$$

$$m = \tau_R + \left(\eta\sqrt{2}\right)\left(\frac{1 - wQ(w)}{Q(w)}\right), \quad (3)$$

where $Q(w) \equiv \sqrt{\pi}\ \mathrm{erfcx}(w)$, and $w \equiv \frac{\eta}{\tau_d\sqrt{2}} + \frac{\tau_R}{\eta\sqrt{2}}$. Since equations (2) and (3) are monotonic, one can invert the equations to find $\eta$ and $\tau_d$ in terms of m and std. Thus, it was possible to change a set of ISIs at one CV to a related set of ISIs at another CV. The cumulative distribution function corresponding to the event probability distribution function given by equation (1) above is:

$$\mathrm{cdf}(t) = 1 - \frac{\mathrm{erfc}(w + (t - \tau_R)\,/\,(\eta\sqrt{2}))}{\mathrm{erfc}(w)}. \quad (4)$$

We randomly assigned numbers drawn uniformly on the unit interval to ISIs in the following way. For each random number, r, we inverted equation (4) to find a $t$ that represented an ISI affiliated with a given mean and CV. By specifying a different mean and CV (and therefore different $\eta$ and $\tau_d$, or equivalently, different $\eta$ and $w$) in equation (4) we derived related sets of ISIs, using a single set of random numbers.

**Analyzing electrophysiology data**

We measured evoked PSP (or FP) peak amplitudes during the last 5 min of pre- and post-conditioning (using pCLAMP) and calculated a post/pre ratio. We evaluated the effects of mTBI, as well as stimulus frequency, temporal pattern, and continuity on the induction of synaptic plasticity [which we defined as: LTD, a significant ($p < 0.05$, *t-test*) decrease (≥ 10%) in post/pre ratios; LTP, a significant increase (≥ 10%) in post/pre ratios; or no change (NC), post/pre ratios that changed < 10% and/or were not significantly different]. For these same time periods (the last 5 min of pre- and post-conditioning), we used pCLAMP to calculate the average PSP: half width (at half height), rise time (10-90%) and decay time (90-10%); as well as the membrane potential and input resistance for each cell.

We also analyzed the kinetic profiles of conditioning responses for individual recordings in the groups that showed a significant difference between control and mTBI. For continuous stimulation paradigms, we took all data points generated during the conditioning period and calculated an exponential and linear best fit, from which we determined the slope. For discontinuous stimulation paradigms, best fits were calculated for each of the nine conditioning sub-epochs and the slopes averaged. Further, we calculated the mean of all response amplitudes (average) as well as the average of the initial 10% (initial) and final 10% (final). For all recordings within a given stimulation paradigm, a simple linear regression analysis was performed to determine the degree to which each of these factors (conditioning period: exponential slope, linear slope, average amplitude, initial amplitude and final amplitude) were correlated with plasticity outcome (post/pre ratio).

#### Calcium imaging

In a subset of the WC patch experiments, the ratiometric calcium sensitive dye Fura-4F (100 μM, Thermo Fisher Scientific Inc.) was added to the pipette solution. At this concentration, the high dissociation constant of the dye ensures it does not alter baseline electrophysiological properties, baseline synaptic transmission, or the induction of synaptic plasticity.[8,9] Calcium imaging experiments were conducted using established methods.[9] Briefly, following a successful WC patch the dye was allowed to equilibrate between the pipette and cytoplasm for at least 20 min. Fluorescent images were then acquired in rapid succession in pairs at the excitation maxima for the calcium-free (336 nm) and calcium-bound (366 nm) forms of Fura-4F using an ultra-high speed wavelength switching xenon arc lamp (Lambda DG-4 system, Sutter Instruments). Acquisition of calcium imaging data and triggering of afferent stimulation were synchronized by using pCLAMP (Axon Instruments, Molecular Devices) to trigger both a signal distribution box (SVB-1, Zeiss) and a pulse stimulator (Master 9, A.M.P.I.). Images were captured using an AxioCam HSm camera mounted on an Axio Examiner D1 microscope equipped with a W plan Apochromat 40x water immersion lens (Zeiss). Fluorescence intensity was measured using AxioVision software (Zeiss).

Calcium calibration was performed using solutions with known calcium concentrations (calcium calibration buffer kit with magnesium #2, Molecular Probes) and 100 μM Fura-4F contained in a synthetic fused silica rectangular capillary (300 μm outer diameter and 50 μm inner diameter, Polymicro Technologies). Calcium dose-response curves were created using the fluorescence intensity data from the calcium-free and calcium-bound forms of Fura-4F as determined by a third-order polynomial interpolation of the data at the excitation wavelengths corresponding to peak fluorescence. A dissociation constant (Kd) was calculated from a fit of the data obtained at either wavelength to a first order Langmur equation. Free cytosolic calcium concentrations were calculated from the background subtracted average fluorescence intensities by the ratio method.[10] We calculated Kd, Rmax and Rmin values by fitting the experimentally determined ratio dose-response curve to the first order Langmur equation. Finally, for each cell the baseline somatic calcium concentration was calculated as an average of 30 measurements made (once every 10 s) during the last 5 min of the pre-conditioning period.

**Histology**

After successful WC recordings, visual cortical slices were processed to verify pyramidal cell morphology and the L2/3 location of recorded cells. Slices were fixed in 4% paraformaldehyde in 0.1M phosphate buffer at 4ºC, then incubated overnight in an avidin biotin horseradish peroxidase complex at 4ºC with 0.1% Triton X-100 (Vectastain Elite ABC Kit, Vector Laboratories). The tissue was counterstained with cresyl violet and cover slipped with dibutyl phithalate (DPX, Electron Microscopy Sciences). Slides were visualized using a Zeiss Axio Imager M2 microscope with a 40x oil immersion objective (EC PlanNeo-Fluar, Zeiss).

**Statistics**

We used SigmaPlot (Systat Software) to determine statistical significance except where otherwise noted. Normality was assessed with a Shapiro-Wilk test. For normally distributed data, we used a *t*-test, or a one-way analysis of variance (ANOVA) followed by Fisher's least significant difference method for multiple comparisons (Fischer's LSD). For datasets with non-normal distributions, we used a Mann-Whitney U test (MWU), or a Kruskal-Wallis one-way ANOVA on ranks (Kruskal-Wallis) followed by Dunn's method for multiple comparisons with Sidak correction (Dunn's corrected; statskingdom.com/kruskal-wallis-calculator.html). Group data are presented as the mean ± standard error (SEM). To assess whether specific characteristics of conditioning responses were predictive of plasticity outcome, for each factor, we calculated a correlation coefficient and determined significance using linear regression. We considered differences statistically significant at $p < 0.05$ (except for Dunn's corrected which was considered significant at $p < 0.017$).
